# Unconjugated bilirubin induces neuro-inflammation in an induced pluripotent stem cell-derived cortical organoid model of Crigler Najjar Syndrome

**DOI:** 10.1101/2023.07.12.548684

**Authors:** Abida Islam Pranty, Wasco Wruck, James Adjaye

## Abstract

Bilirubin-induced neurological damage (BIND), which is also known as Kernicterus, occurs as a consequence of defects in the bilirubin conjugation machinery, thus resulting in unconjugated bilirubin (UCB) to cross the blood–brain barrier (BBB) and accumulation. Severe hyperbilirubinemia can be caused by a mutation within the *UGT1A1* encoding gene. This mutation has a direct contribution towards bilirubin conjugation leading to Kernicterus as a symptom of Crigler Najjar Syndromes (CNS1, CNS2) and Gilbert syndrome, which results in permanent neurological sequelae. In this comparative study, we used human induced pluripotent stem cells (hiPSCs)-derived 3D-brain organoids to model BIND *in vitro* and unveil the molecular basis of the detrimental effects of UCB in the developing human brain. hiPSC-derived from healthy and CNS patients were differentiated into day-20 brain organoids, these were then stimulated with 200nM UCB. Analyses at 24- and 72-hrs post-treatment point at UCB-induced neuro-inflammation in both cell lines. Transcriptome and associated KEGG and Gene Ontology analyses unveiled activation of distinct inflammatory pathways such as cytokine-cytokine receptor interaction, MAPK signaling, calcium signaling, NFκB activation. Furthermore, both mRNA expression and secretome analysis confirmed an upregulation of pro-inflammatory cytokines such as IL6 and IL8 upon UCB stimulation. In summary, this novel study has provided insights into how a human iPSC-derived 3D-brain organoid model can serve as a prospective platform for studying the etiology of BIND-Kernicterus.

## 1. Introduction

Clinical jaundice is a commonly occurring transitional condition, caused by unconjugated hyperbilirubinemia, which affects about 85% of newborns in their first week of postnatal life [1,2]. Unconjugated hyperbilirubinemia is normally benign, nevertheless, the protective mechanisms of the brain can be surpassed in the infants when the unconjugated bilirubin (UCB) starts accumulating and cross the blood-brain barrier due to a defective bilirubin conjugation machinery. Elevated UCB levels result in severe brain injury including short-term and long-term neurodevelopmental disabilities, which can progress into acute or chronic bilirubin encephalopathy, known as kernicterus or bilirubin-induced neurological dysfunction (BIND) [3]. A genetic-based disorder named Crigler–Najjar Syndrome (CNS) may also cause BIND. Bilirubin-UGT1 (UGT1A1) is the only isoform of the uridine-diphosphoglucuronate glucuronosyltransferases (UGTs) enzyme family that specifically contributes to bilirubin conjugation. Therefore, a mutation in the gene encoding UGT1A1 can cause CNS type 1 and 2 by complete or partial inactivation of the enzyme, which prevents the liver from metabolizing bilirubin [4]. This hinders bilirubin conjugation, causing UCB to accumulate in serum, cross the blood–brain barrier, and eventually deposited in the basal ganglia or cerebellum, thereby resulting in BIND [5-7]. The mechanism underlying BIND and the correlation between UCB level and neurological abnormalities are not well understood.

Neuro-inflammation is one of the key features observed in distinct model systems of bilirubin induced brain toxicity. It has been shown that in *UGT1A1* knocked-out mice Toll-like receptor 2 (TLR2) is required for regulating gliosis, pro-inflammatory mediators, and oxidative stress when neonatal mice are exposed to severe hyperbilirubinemia but TLR2 also has anti-apoptotic property. This correlates with the upregulation of TNF-α, IL-1β, and IL-6, thus indicative of neuro-inflammation in the CNS [8]. TNF-α and NFκB are the key mediators of bilirubin-induced inflammatory responses in murine models [9]. Sustained exposure of developing mouse cerebellum with UCB induces activation of oxidative stress, endoplasmic reticulum (ER) stress and inflammatory markers, thus pointing to inflammation as a key contributor of bilirubin-induced damage in conjunction with ER stress in the onset of neurotoxicity [9]. Immature rat neurons also manifested distinct features of oxidative stress and cell dysfunction upon UCB exposure by increased ROS production, disruption of glutathione redox status and cell death [10]. Bilirubin induced apoptosis and necrosis-like cell death leading to neuritic atrophy and astrocyte activation have also been observed in monotypic nerve cell cultures [11].

Elevated UCB levels perturb plasma, mitochondrial, and/or ER membranes of neuronal cells, probably leading to neuronal excito-toxicity, mitochondrial energy failure, or increased intracellular calcium concentration [Ca^2+^], which are assumed to be linked to the pathogenesis of BIND. Increased [Ca^2+^] and the following downstream events may activate proteolytic enzymes, apoptotic pathways, and/or necrosis, depending on the intensity and duration of bilirubin exposure [12].

In this study, we used hiPSC-derived 3D-brain organoids as a potential *in vitro* model system to enhance our meagre understanding of BIND-associated molecular pathogenesis and the breadth of complexities at the cellular and molecular levels that accompany the detrimental effects of UCB to the developing human brain. hiPSC derived from a healthy and CNS patient were differentiated into day-20 brain organoids, and then stimulated with 200nM UCB. We performed further analyses and observed the induction of neuro-inflammation along with activation of cytokine-cytokine receptor interaction, calcium signaling pathway, MAPK signaling pathway, and neuroactive ligand receptor interaction as processes leading to BIND.

## 2. Methods and materials

### 2.1. Cell cultivation and formation of neural cortical organoids

IPSC lines from a healthy individual-UM51 and CNS patient-i705-C2 were used in this study [13,14]. Cells were plated on Matrigel (Corning, New York, NY, USA) coated culture dishes using mTeSR plus medium (StemCell Technologies, Vancouver, Canada). Cultures were routinely tested for mycoplasma contamination. Cells were dissociated into small aggregates with ReLeSR (StemCell Technologies) every 5-7 days and split in a 1:5 ratio into fresh Matrigel-coated dishes. Alternatively, the cells were also split as single cells using accutase (Life Technologies, Waltham, MA, USA) while seeding for organoid generation. The protocol described by Gabriel et al.2016, was employed to differentiate iPSCs into cortical organoids with minor modifications [15]. Briefly, 20,000 single iPSCs were seeded onto each well of a U-botton 96-well plate (NucleonTM SpheraTM, Thermo Fisher Scientific, Rockford, IL, USA) to form embroid bodies with mTesR plus medium and 10μM ROCK inhibitor Y-27632 (Tocris Bioscience, Bristol, UK). The EBs were cultured in the plate for 5 days with neural induction medium (Stem cell technologies) to initiate neural induction. On day-6, the EBs were transferred into a bioreactor (PFIEFFER) with differentiation medium consisting of DMEM/F12 and Neural Basal Medium (in 1:1 ratio), supplemented with 1:200 N2, 1:100 L-glutamine, 1:100 B27 w/o vitamin A, 100 U/ml penicillin, 100 mg/ml streptomycin, 0.05 mM MEM non-essential amino acids (NAA), 0.05 mM ß-mercaptoethanol (all from Gibco, Waltham, MA, USA), and 23 μM insulin (Sigma, Taufkirchen, Germany) (see Table S1). The spinner flasks were coated with anti-adherenet rinsing solution (StemCell technologies) before transferring the EBs. Transferred EBs were counted as day-0 organoids from this time point (differentiation day-6), as the spheres were transferred into the spinner flask to spontaneously patternized into cortical organoids. From day-9 onwards, 0.5 μM of dorsomorphin (Tocris Bioscience) and 5μM SB431542 (Tocris Bioscience) were added in addition to the differentiation medium and the medium in the bioreactor was changed once a week. HLCs were derived from iPSCs following the protocol described by Graffmann et al.2016 (Supplementary figure S1) [16].

### 2.2. Cryosectioning

Cells (HLCs) were fixed in 4% paraformaldehyde (PFA) (Polysciences, Warrington, FL, USA) for 10 min and cortical organoids were fixed for 30 min at 37°C. After washing with PBS, the cells were directly used for staining and the organoids were dehydrated in 30% sucrose in PBS overnight at 4 °C. Then the organoids were embedded using Tissue-Tek OCT Compound (embedding medium) (Sakura Finetek, Umkirch, Germany) in cryo-molds and snap-frozen in 2-methylbutan (Carl Roth, Karlsruhe, Germany) and dry ice. The embedded organoids were stored at -80°C. Organoids were sectioned into 15 μm sections using a Cryostat (CM1850, Leica, Nussloch, Germany) and captured in Superfrost plus slides (Thermo Scientific). The sectioned organoid slices were stored on -80°C prior to immunofluorescnece.

### 2.3. Immunocytochemistry

Fixed cells were permeabilized for 10 minutes with 0.1% TritonX in PBS+Glycine (30mM Glycine) at RT. They were then washed once with PBS and then the unspecific binding sites were blocked for 2hr at RT with blocking buffer 0.3% BSA in PBS+Glycine. Frozen sections were thawed at RT and PBS was added drop by drop without touching the organoid sections and incubated at RT for 15 minutes. Tissue Tek was washed off with PBS and the slide was washed once more with PBS. The sections were permeabilized with 0.7% TritonX100 + 0.2% Tween20 in PBS+Glycine for 15min at RT. After permeabilization blocking was carried out for 2 hrs at RT with 0.2% TritonX100 + 0.3% BSA in PBS+Glycine. For the cells and organoid sections, the primary antibody solution was incubated overnight at 4 °C (see Table S2). After removing the primary antibodies and thorough washing, the secondary antibodies were added for 2 hrs and incubated at RT. Nuclei were stained with Hoechst. Stained cells and sections were imaged using a Zeiss fluorescence microscope (LSM 700). Particular staining regions were observed under a Zeiss confocal microscope (LSM 700). Individual channel images were processed and merged with ImageJ.

### 2.4. TUNEL assay

Apoptotic cells were detected using the DeadEnd™ Fluorometric TUNEL System (Promega, G3250, USA) following the manufacturer′s protocol.

### 2.5. Reverse Transcriptase PCR (RT-PCR)

The cells (HLCs) and cortical organoids were lysed in Trizol to isolate the RNA. 7-8 UCB treated and non-treated organoids were taken for RNA isolation. RNA was isolated with the Direct-zol™ RNA Isolation Kit (Zymo Research) according to the user’s manual including 15 minutes and 30 min DNase digestion step for cells and organoids respectively. 500 ng of RNA were reverse transcribed using the TaqMan Reverse Transcription Kit (Applied Biosystems). Primer sequences are shown in Table S3. Real time PCR was performed in technical and independent experiment triplicates with Power Sybr Green Master Mix (Life Technologies) on a VIIA7 (Life Technologies) machine. Mean values were normalized to RPL0 and fold-change was calculated using the indicated controls. The observed fold changes are depicted as mean values with 95% confidence interval (CI).

### 2.6. Human XL Cytokine Assay

The conditioned medium or supernatant of control and UCB treated cortical organoids from both 24 and 72 hrs treatments were stored and used for the proteome profiler antibody array. Relative expression level of 105 soluble human proteins and cytokines were determined using the Human XL Cytokine Array Kit from R&D Systems. The cytokine array was performed following the manufacturer’s guidelines. In brief, membranes were blocked for 1 hr on a rocking platform using the provided blocking buffer and then the samples were prepared by diluting the desired quantity to a final volume of 1.5 ml with distinct array buffer (array buffer 6). The sample mixtures were pipetted onto the blocked membranes and were incubated overnight at 4°C on a rocking platform. Membranes were then washed three times with washing buffer for 10 minutes each at RT. Then the membranes were incubated with the Detection Antibody Cocktail for 1 hr at room temperature and then wash three times thoroughly. Afterwards Streptavidin-HRP was added onto the membranes, which were incubated for 30 min at room temperature. ECL detection reagent (Cytiva, Freiburg, Germany) was used to visualize the spots on the membrane and then detected in a Fusion FX instrument (PeqLab).

### 2.7. Image and Data Analysis of the Human XLCytokine Array

After performing the cytokine array with nontreated and UCB treated organoids, hybridizations of the cytokine assays were scanned and read into the FIJI/ImageJ software, where spots were quantified as described in our previous publication by Wruck et al. [17,18]. Spots were associated with cytokine identifiers provided by the manufacturer (Proteome Profiler Array from R & D Systems, Human XL Cytokine Array Kit, Catalog Number ARY022B). Integrated densities of the spots as result of the quantification were read into the R/Bioconductor [19]. The Robust Spline Normalization from the R/Bioconductor package lumi was applied to the data [20]. Expressed and differentially expressed cytokines were determined as described before [17].

### 2.8. Analysis of Gene Expression Data

i705-C2 and UM51 organoid samples, untreated and treated with UCB were measured after 24 and 72 hrs on the Affymetrix Human Clariom S Array at the core facility of the Heinrich-Heine Universität, Düsseldorf (BMFZ: Biomedizinisches Forschungszentrum). Processing of the microarray data was performed in the R/Bioconductor environment [19]. The background correction and normalization with the Robust Multi-array Average (RMA) method was achieved by the Bioconductor package oligo [21]. Using values of dedicated background spots on the microarray a statistic was calculated to determine a detection-p-value to judge if a probeset is expressed (detection-p < 0.05) as was described before in Graffmann et al. [16]. Probesets expressed following this criterion were mapped to unique gene symbols according to annotations provided by Affymetrix and compared in venn diagrams via the VennDiagram package[22]. The function heatmap.2 from the R gplots package was applied to draw heatmaps and associated clustering dendrograms using Pearson correlation as similarity measure and color-scaling by Z-scores of rows (genes) [23]. Genes with a detection-p-value below 0.05 in both conditions were considered up-regulated when the ratio was greater than 1.5 or down-regulated when the ratio was less than 0.67.

### 2.9. Analysis of Pathways and Gene Ontologies (GOs)

For the analysis of KEGG (Kyoto Encyclopedia of Genes and Genomes) pathways association between genes and pathways had been downloaded from the KEGG website in July 2020 [24]. Over-representation of pathways in gene sets of interest was tested via the R-builtin hypergeometric test. The R package GOstats was applied for calculating over-represented gene ontologies [25]. For dot plots of the most significantly over-represented pathways the R package ggplot2 was employed [26].

### 2.10. Western Blotting

Total protein from cortical organoids and HLCs were isolated using RIPA buffer (Sigma-Aldrich) consisting of protease and phosphatase inhibitors (Roche, Mannheim, Germany). Afterwards the Pierce BCA Protein Assay Kit (Thermo Fisher) was used to determine the Protein concentration. Approximately 20μg of heat denatured protein lysate of each sample was loaded on a 4-12% SDS-PAGE and then transferred by wet blotting to a 0.45 μm nitrocellulose membrane (GE healthcare). After 1 hr of blocking with 5% Milk in TBST, the membranes were stained with anti-P53, anti-γH2A.X, anti-UGT1A1, anti-CREB, anti-phospho-CREB, anti-phospho-P38MAPK antibodies. Incubation with primary antibodies was performed overnight at 4°C. After washing the membranes three times with TBST, the secondary antibody incubation was performed for 2 hrs at RT followed by washing with TBST (Table, S2). Anti ß-actin and anti-GAPDH were used as housekeeping proteins to normalize protein expression. ECL Western Blotting Detection Reagents (Cytiva, Freiburg, Germany) was used to visualize the stained protein bands and then detected in a Fusion FX instrument (PeqLab). Band intensity quantification and analysis was performed with Fusion Capt Advance software (PeqLab) and was normalized to beta-actin band intensity.

### 2.11. Measurement of cytochrome P450 activity

The P450-Glo™ CYP2D6 Assay and P450-Glo™ CYP3A4 Assay Luciferin-IPA (Promega) kits were used to measure Cytochrome P450 2D6 and P450 3A4 activity employing a luminometer (Lumat LB 9507, Berthold Technologies).

## 3. Results

### 3.1. iPSC-derived cortical organoids show typical cortical neuronal features

hiPSCs derived from renal progenitor cells isolated from urine of a 51year old healthy male of African origin (UM51) and fibroblasts cells from a Crigler-Najjar Syndrome (CNS) male patient (i705-C2) were cultured as colonies and used to generate 3D-cortical neuronal organoids in triplicates. i705-C2 cells retained the *UGT1A1* mutation, as the cells were derived from Crigler-Najjar syndrome (CNS) patient (Supplementary figure S1) [13,14]. The iPSCs were seeded as single cells to form EBs and then transferred into a bioreactor to grow spontaneously as organoids (Figure 1a, 1b). Similar to previously established cerebral organoids, our generated organoids recapitulate human cortical developmental features with progenitor zone organization and the presence of radial glia stem cells and cortical neurons (Figure 1c, 1d) [27]. The self-patterned organoids showed cortical neuronal identity with the expression of the radial glia marker-SRY-box transcription factor 2 (SOX2), neuronal markers Beta III Tubulin (TUJ1), microtubule associated protein 2 (MAP2), and Doublecortin (DCX) at day-15 (Figure 1c, 1d).

**Figure 1:**
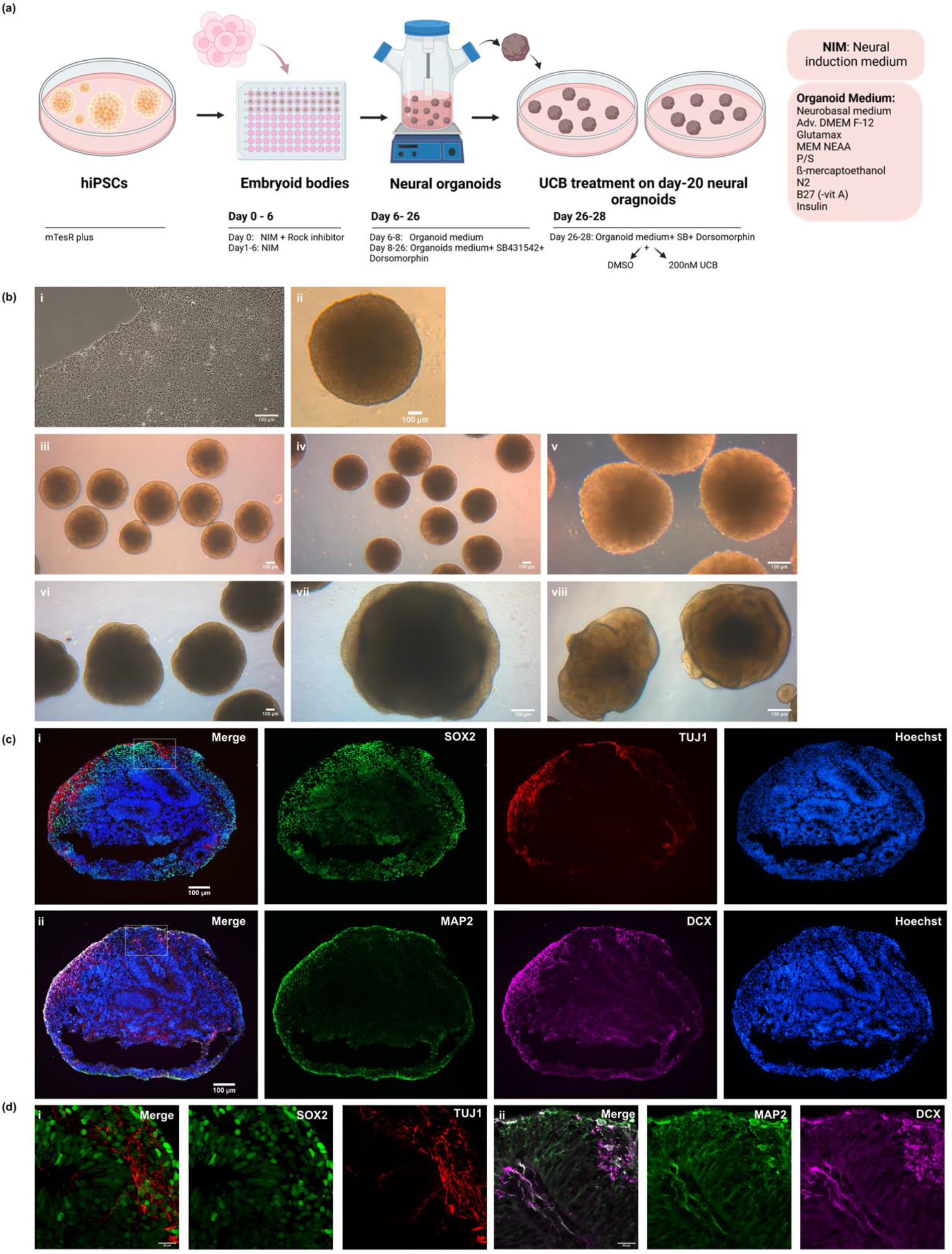
Generation of iPSC-derived cortical brain organoids. **(a)** Schematic depiction of the generation of cortical organoids **(b)** Bright-field images showing the cortical brain organoid generation from iPSCs to organoids. Scale bars depict 100μm (i) iPSCs culture from patient derived iPSC line (i705-C2) (ii). EBs were formed after 24 hrs of seeding in u-bottom 96 well plates. (iii, iv, v) Day-1 organoids (day-7 of differentiation). (vi. vii. viii) Day-20 organoids before UCB treatment. **(c)** Neural identity of day-15 cortical organoids was confirmed by the expression of (i) the radial glia marker SOX2 (green) and neuronal marker TUJ1 (red), (ii) MAP2 (green) and DCX (magenta), on sections and IF staining. Scale bars depict 100 μm. **(d)** Confocal imaging of the SOX2 (green), TUJ1 (red) (i), MAP2 (green) and DCX (magenta) (ii) showing a higher magnification. Scale bars depict 20μm.

### 3.2. UCB induced neuro-inflammation with elevated expression of pro-inflammatory cytokines

Day-20 cortical organoids were treated continuously with lipophilic unconjugated bilirubin (UCB) for 72 hrs in the bioreactor (Figure 1a). RNA from UM51 and i705-C2 cortical organoids were isolated in triplicate at 24 hrs and 72 hrs time point of UCB treated and control conditions to investigate expression of pro-inflammatory-associated cytokines at the mRNA level. In parallel, the supernatant was collected at both time points to carry out further secretome analysis using cytokine arrays (Supplementary Figure S2b). qRT-PCR analysis from the i705-C2 line showed 3-fold increase in *IL6* and *IL8* expression at 24 hrs post-treatment, but decreased to 2-fold at 72 hrs, while *IL8* gene expression was 3-fold enhanced also for the UM51 healthy control line (Figure 2a). *IL6* mRNA expression was 1.5-fold enhanced only at 72 hrs post-treatment in the healthy line. On the other hand, expression of *TNF-α* did not show any change upon UCB exposure for the UM51 line, but a slight increase with 1.5-fold (24 hrs and 72 hrs) change was observed in the i705-C2 line. Additionally, a heatmap obtained from the cytokine array-based analysis showed the secretome profile of various cytokines after UCB treatment for both cell lines at 24 hrs and 72 hrs (Figure 2b) (Supplementary figure S2b). The enhanced secretion of the pro-inflammatory cytokines such as IL6, IL8, TNF-α, IL1-ß, IL16, IL1-ra, IL-11, INF-G, was observed along with anti-inflammatory cytokine IL-13 secretion (Figure 2b). Secretome analysis also revealed enhanced vascular endothelial growth factor (VEGF) secretion and reduced Sex hormone-binding globulin (SHBG) secretion upon UCB treatment for all the conditions, except UM51 24hrs (Figure 2b) (Supplementary figure S2b). Healthy line showed 2-fold increase of *VEGF* mRNA expression at 72 hrs and patient line showed 1.4-fold and 1.6-fold increase at 24 and 72 hrs respectively (Supplementary figure S2c). On the other hand, *SHBG* expression was slightly decreased with 0.6-fold (UM51) and 0.8-fold (i705-C2) at 72 hrs post-treatment (Supplementary figure S2c). Overall, the increase in both mRNA expression and protein secretion of *IL6, IL8 and TNF-α* implies the initiation of neuro-inflammation in the cortical organoids upon UCB treatment.

**Figure 2:**
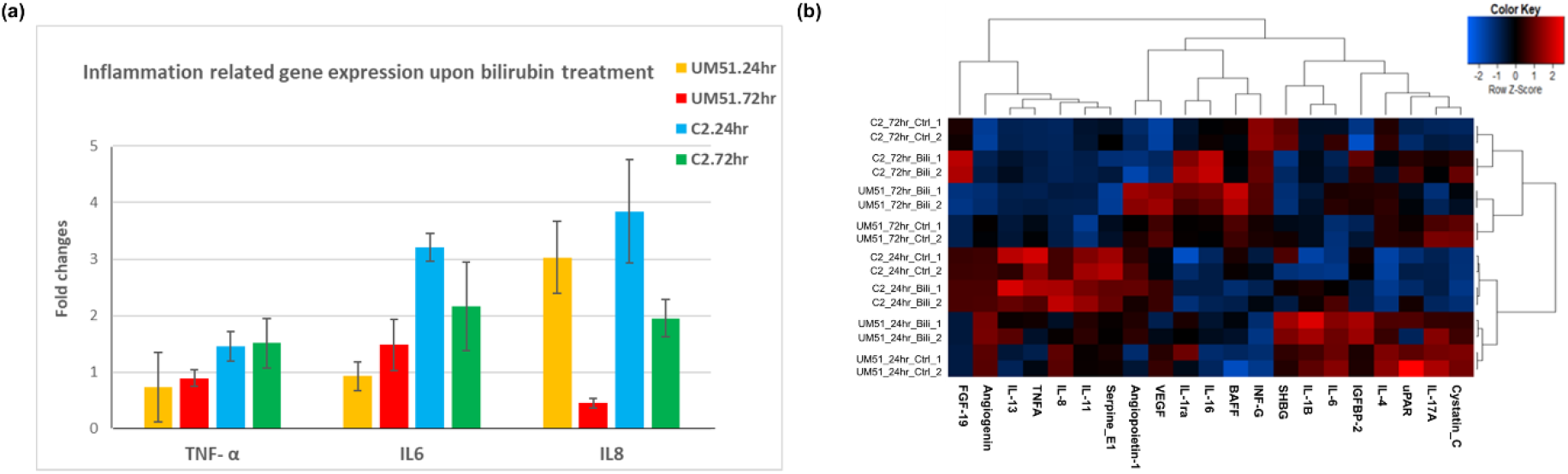
Pro-inflammatory cytokine expressions are increased by UCB. **(a)** qPCR analysis showing an increased mRNA expression of pro-inflammatory cytokines such as *IL6* and *IL8* for both cell lines, while *TNF-α* was upregulated only for the patient line. Inflammatory responses were observed at 24 hrs in the patient line but declined gradually at 72 hrs, whereas the initiation of inflammatory responses shifted at 72 hrs for the UM51 line. Depicted values are mean of three different independent (n=3) experiments. Error bar depicts ± 95% confidence interval. Values were normalized to RPL0 (housekeeping gene) and subsequently to DMSO treated control organoids. **(b)** Cytokine array-based secretome analysis showed increased IL6, IL8 and TNF-α secretion along with other cytokines upon UCB exposure, while the healthy control line showed lower IL6, IL8 and TNF-α secretion compared to the patient line.

### 3.3. Distinct inflammation-associated pathways are activated by UCB in cortical organoids

To further investigate the neuro-inflammatory effect of UCB on cortical organoids at the molecular level, RNA was isolated from 24 and 72 hrs post-treated and untreated (control) UM51 and i705-C2 cortical organoids for Microarray analysis. Venn diagrams obtained from this transcriptome analysis showed that the i705-C2 cortical organoids expressed 14721 genes in common between treatment and control and uniquely expressed 906 genes at 24 hrs post-treatment, whereas at 72 hrs, 15317 genes were expressed in common, and 124 genes were uniquely expressed in i705-C2 organoids (Figure 3a). On the other hand, UM51 cortical organoids expressed 15491 and 15731 common set of genes, while uniquely expressed 289 and 351 genes upon UCB exposure at 24 hrs and 72 hrs respectively (Figure 3a). KEGG pathways associated with the uniquely expressed genes at 24 hrs post treatment included, *TNFSF12, AZI2* and for example cytokine-cytokine receptor interaction was activated in the i705-C2 line, which gradually decreased at 72 hrs (Figure 3b, 3c) (Supplementary figure 3b). The cytokine-cytokine receptor activation was not observed in the healthy UM51 line at 24 hrs but at 72 hrs (Figure 3d) (Supplementary figure S3c). However, Calcium and MAPK signaling pathways, and neuro-active ligand receptor interactions were activated at 24 hrs for the patient line and for both cell lines at 72hrs post UCB exposure (Figure 3b, 3c, 3d) (Supplementary figure S3a).

**Figure 3:**
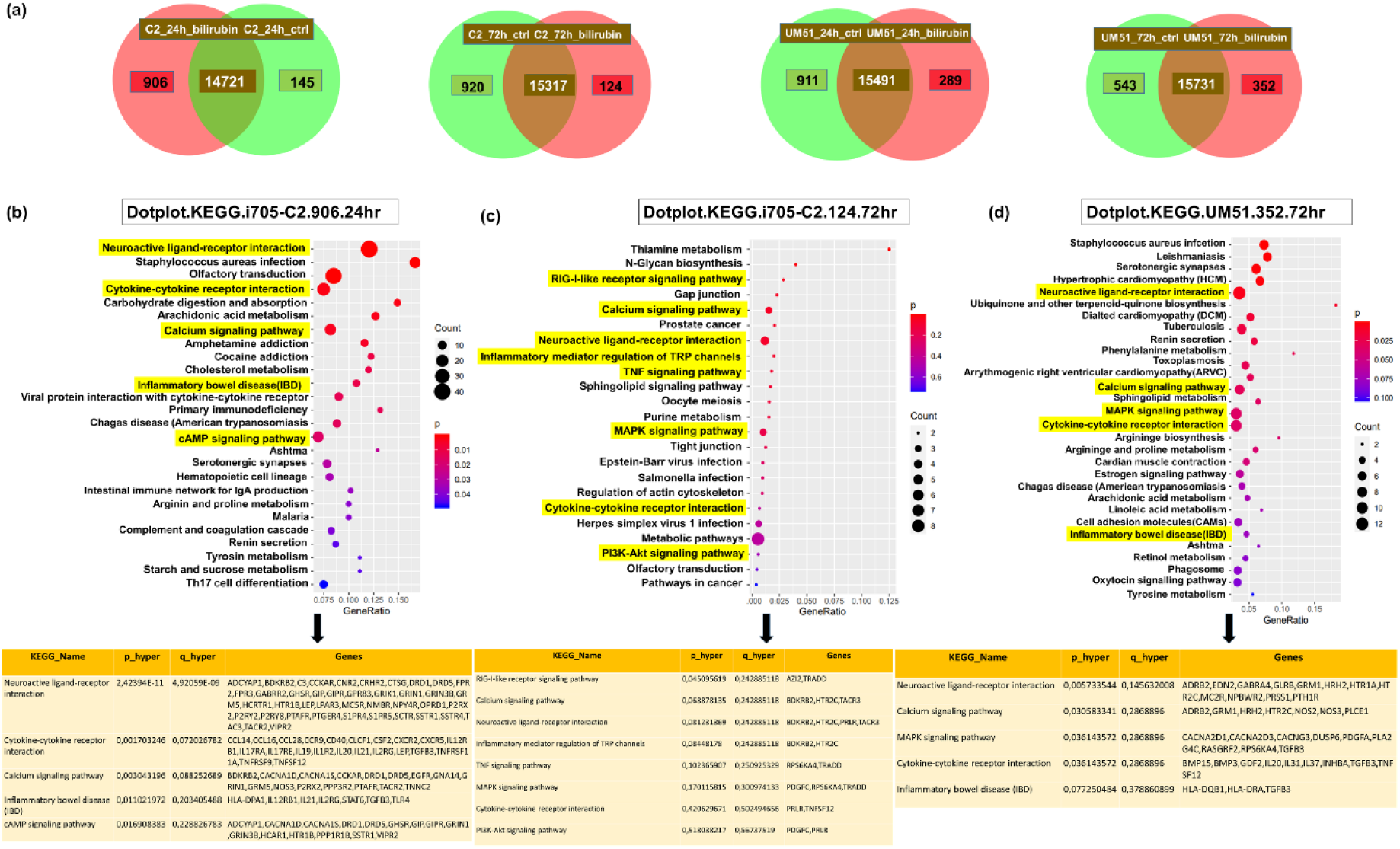
Overview of distinct activated pathways upon UCB treatment in cortical brain organoids. **(a)**. Venn diagrams show the uniquely and differentially expressed genes upon bilirubin treatment for UM51 and i705-C2 cell line for both the 24 hrs and 72 hrs time points. Dot plots from KEGG associated pathways and corresponding genes reveal cytokine-cytokine receptor activation in the patient line at 24 hrs post-treatment and **(b)**, which goes gradually down at 72 hrs **(c)**. This cytokine-cytokine receptor activation was not observed in UM51 line at 24 hrs (Supplementary figure 3c) but observed at 72 hrs **(d)**. Dot plots show the activation of calcium signaling pathway, MAPK signaling pathway, and neuro-active ligand receptor interaction for both cell lines at 72 hrs post UCB exposure.

### 3.4. UCB treatment differentially regulate DNA damage and repair related pathways

Transcriptome analysis of the cortical organoids provided an overview of the GO terms and associated KEGG pathways of the differentially regulated genes (common set of genes) upon UCB exposure. KEGG pathways revealed DNA damage and repair-related pathways such as P53 signaling pathway, homologous recombination, Fanconi Anemia pathway were upregulated in the UM51 line and downregulated in the i705-C2 line at 24 hrs (Figure 4b ii, iii). In parallel, at 24 hrs the i705-C2 line showed activation of NFκB, PI3K, and chemokine signaling pathways indicating the initiation of inflammation, while the cellular developmental processes related to Notch and TGFß signaling pathways were upregulated (Figure 4b i). Interestingly, a number of these inflammatory and development related pathways (neuroactive-ligand receptor interaction, cAMP signaling pathway, cytokine-cytokine receptor interaction) were observed to be downregulated at 24 hrs in the UM51 line (Figure 4B iv). The mRNA expression of *NLRP3* was enhanced only for the i705-C2 line, which is an upstream activator of NFKß signaling and plays a role in inflammation, immune response, and apoptosis (Supplementary figure S3b). UM51 showed slight increase in both mRNA (1.2-fold) and protein expression of CREB (1.3-fold) and protein expression for phospho-CREB (1.13-fold), while the i705-C2 line showed only slightly enhanced mRNA expression (Supplementary figure S3b, S4a, S4b). Moreover, the UM51 line showed slightly enhanced mRNA expression of the DNA damage-repair and apoptosis-related genes at 24 and 72 hrs such as P53 (1.24-fold, 1.14-fold), *BCL2* (1.5-fold, 1.3-fold), *ATM* (1.41-fold), *ATR* (1.3-fold, 1.3-fold), *CHEK1*(1.3-fold, 1.2-fold), and *CHEK2* (1.6-fold) compared to the i705-C2 line (Figure 4a i). However, the patient line showed increased *BCL2* (1.3-fold) and *MDM2* (1.4-fold) expression at 24hrs and increased *P53* (1.6-fold) expression at 72 hrs post UCB treatment. In addition to that, WB analysis revealed a 1.64-fold increase in P53 protein expression at 24 hrs in the healthy line and with a non-significant 1.1-fold increase at 72 hrs in the patient line (Figure 4a ii). Furthermore, GO-terms of the common set of expressed genes revealed neurodevelopmental pathways to be differentially upregulated with the activation of cellular developmental process, nervous system development, axon development, axon guidance, tight junction, positive regulation of axogenesis, positive regulation of synapse assembly for both cell lines at 72 hrs post-UCB treatment (Supplementary figure S4d).

**Figure 4:**
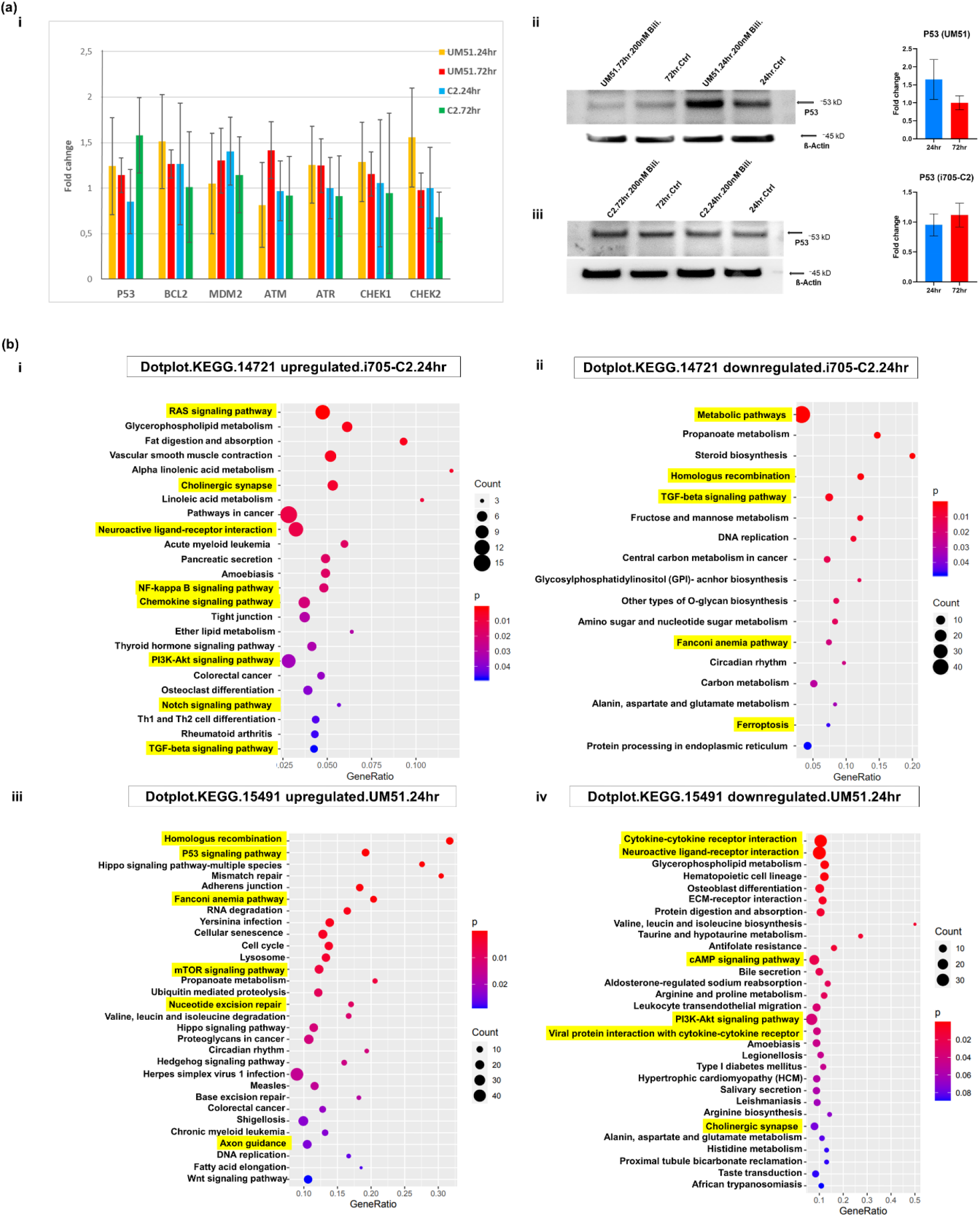
UCB treatment differentially regulated DNA damage and repair related pathways. **(a)** (i) qRT-PCR analysis shows DNA damage and repair related gene (*P53, BCl2, MDM2, ATM, ATR, CHEK1, CHEK2*) expression after UCB treatment. Depicted values are mean of three different independent (n=3) experiments. Error bar depicts ± 95% confidence interval. Values were normalized to RPL0 (housekeeping gene) and subsequently to DMSO treated control organoids. (ii, iii) WB analysis shows increased P53 protein expression at 24 hrs in the healthy line and at 72 hrs in the patient line, thus implying DNA damage and/or apoptotic cell death. Bar graphs showing the protein expression in fold change. Depicted values are mean of three different independent (n=3) experiments. Values were normalized to ß-Actin (A, B) (housekeeping gene) and subsequently to DMSO treated control organoids. **(b)** Dotplots from KEGG pathways reveal that DNA damage and repair-related pathways such as P53 signaling pathway, homologous recombination, Fanconi Anemia pathways are upregulated in the healthy line at 24 hrs (iii) and downregulated in the patient line (ii). Additionally, at 24 hrs the patient line showed activation of NFκB, PI3K, and chemokine signaling pathways thus implying the onset of inflammation (i). Cytokine-cytokine receptor interaction, neuroactive ligand-reactor interaction, cAMP signaling pathway were downregulated in the healthy line at 24 hrs (iv).

### 3.5. UCB induces apoptotic cell death on cortical organoid

Apoptotic cell death in UCB treated cortical organoids was evaluated by immunofluorescence and Western blot analysis. Both cell lines exhibited an increased level of cell death in the presence of terminal deoxynucleotidyl transferase dUTP nick end labeling (TUNEL) at 24 hrs post-treatment (Figure 5a, 5b). Cleaved caspase 3 positive cells were observed in the organoid sections, pointing at apoptotic cell death at 24 hrs post treatment (Figure 5a, 5b).

**Figure 5:**
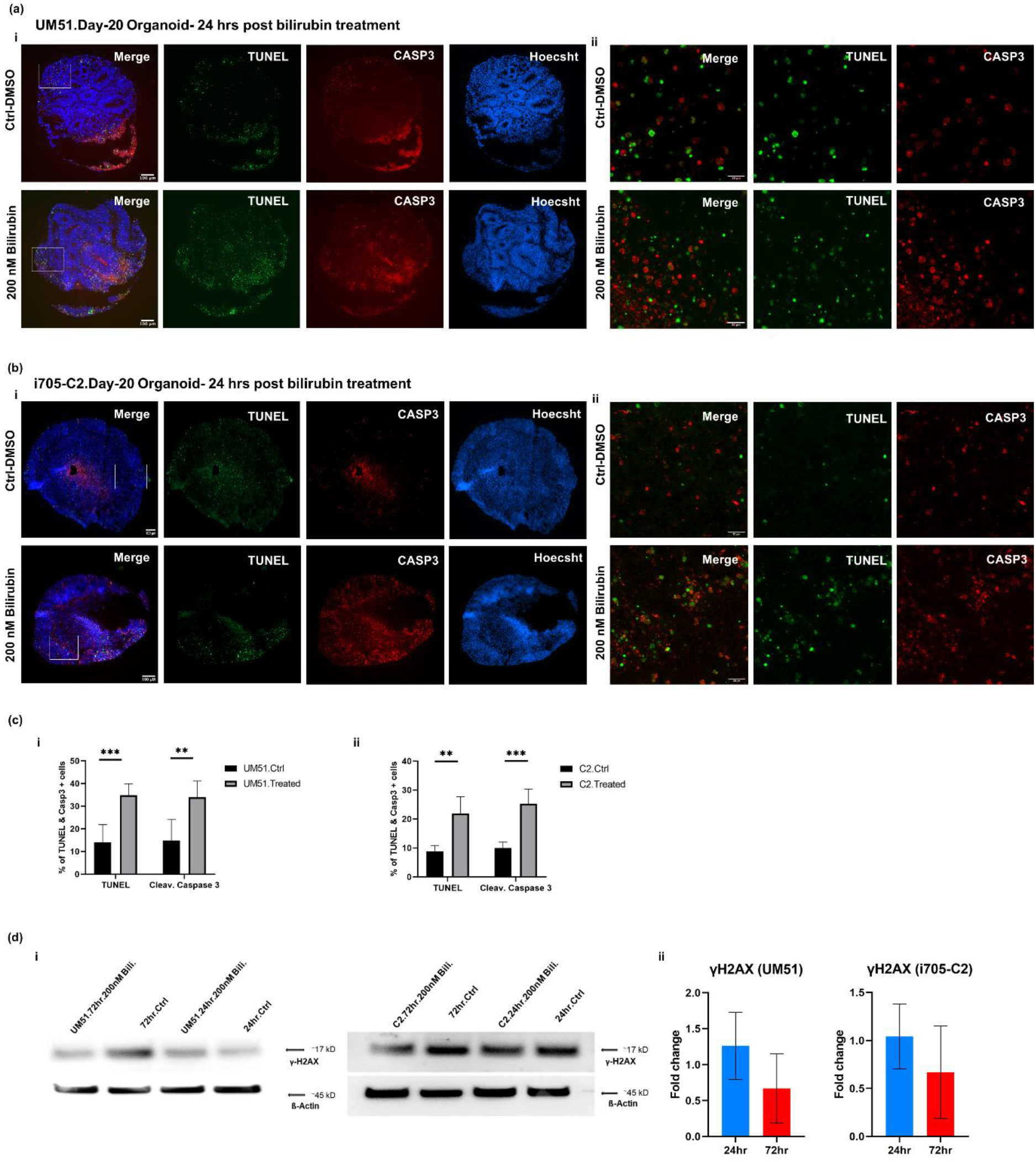
UCB treatment initiates apoptotic cell death on cortical neurons. IF staining of Day-20 cortical organoid sections after 24 hrs of UCB exposure for the healthy line **(a)** and patient line **(b)**. In panels **(a)** (i), **(b)**. (i), **(c)**. (i,ii) we see increased apoptotic cell death as confirmed with the presence of TUNEL positive (green) staining and apoptotic cell death marker cleaved CASP 3 (red). Scale bars depict 100 μm. **(a)** (ii), **(b)**.(ii) Confocal imaging of the TUNEL (green), cleaved CASP 3 (red) positive cells showing a higher magnification. Scale bars depict 20μm. **(c)** (i,ii) Bar graphs indicate quantification of TUNEL and cleaved CASP3 staining (N=6). Asterisk depicts significance, which is determined by α-value ≤ 0.05. **(d)** WB anaylsis showed increased γH2AX expression at 24hrs post UCB treatment (N=3). Bar graphs showing the protein expression in fold change. Values were normalized to ß-Actin (housekeeping gene) and subsequently to DMSO treated control organoids

Immunofluorescence-based quantification revealed a 34% increase in TUNEL and cleaved Caspase 3 positive cells in the UM51 organoid sections, whereas the i705-C2 sections had 22% TUNEL positive and 25% cleaved Caspase 3 positive cells at 24 hrs (Figure 5c). γH2AX expression, which indicates both DNA damage or apoptotic cell death, showed an enhanced protein expression of 1.26-fold based on WB analysis on the healthy line after 24 hrs of UCB exposure, whilst the patient line showed almost no change in γH2AX expression (1.04-fold increase) (Figure 5d). The Immunofluorescence-based staining also showed a slight increase in γH2AX expression at 24 hrs post UCB treatment (Supplementary figure S5). Further indication of DNA damage was observed with the upregulation of P53-mediated signaling pathway in the KEGG analysis of the UM51 transcriptome at 24 hrs (Figure 4b iii).

## 4. Discussion

The postnatal condition-unconjugated hyperbilirubinemia, can lead to kernicterus [3]. Kernicterus is a complex neuropathological ailment, also termed as UCB encephalopathy and can lead to acute or chronic neurological disabilities, resulting as bilirubin-induced neurological dysfunction (BIND) [28,29]. A genetic disorder caused by defective enzyme-UGT1A1 leads to Crigler–Najjar Syndrome (CNS) and can also cause BIND [30]. The complete or partial mutation within the *UGT1A1* gene hinders bilirubin conjugation, which assists lipophilic UCB to cross the blood brain barrier (BBB), resulting in BIND [5,7,31]. In this study, we successfully generated iPSC-derived 3D cortical organoids to model BIND *in vitro* and unveil insights into the detrimental effects of UCB in the developing human brain at the molecular and protein level. Our *in vitro* model is comprised of a healthy iPSC (UM51) and CNS patient derived (i705-C2) iPSC, harboring the *UGT1A1* mutation [13,14]. The absence of UGT1A1 protein was confirmed after differentiation of the iPSCs into hepatocyte-like cells (HLCs). Day-20 cortical organoids were continuously treated with 200 nM UCB for 24 hrs and 72 hrs to observe the possible short-term and long-term UCB-induced effects [3,32].

UCB can interfere BBB integrity by GSH (Glutathione) disruption and increased endothelial nitric oxide synthase (NOS) by enhanced cytokine release [3,33]. Cytokines are one of the major effector groups of the inflammatory cascade. During brain injury, they play a key role in regulating nerve cell responses [34]. Cytokines might have both beneficial and detrimental effects on neurons depending on their secretion balance [28,35,36]. Production of pro-inflammatory mediators can cause neuronal apoptosis and neuro-inflammation [37,38]. Even though, microglia are the key cell type in the CNS which secrete pro-inflammatory cytokines upon stress, UCB treated neurons showed enhanced secretion of IL6 with decreased secretion of IL-1ß [39]. Based on these observations, we analysed mRNA expression of several pro-inflammatory cytokines and also their secretome profile. mRNA expression from control and UCB treated conditions showed 3-fold increase in *IL6* and *IL8* expression at 24 hrs post treatment in the i705-C2 (patient) line which then decreased to 2-fold at 72 hrs. A similar pattern was observed in the secretome analysis for the i705-C2 line with an increased secretion of IL6 and IL8 at 24 hrs post-treatment, then decreased at 72 hrs. This observation might imply that inflammatory responses were initiated earlier in the i705-C2 line and gradually reverted to normal levels with time as a consequence of cellular defense mechanisms, establishing homeostasis. On the other hand, the UM51 (healthy) line showed 3-fold enhanced *IL8* mRNA expression only at 24 hrs, while secretion of IL8 protein was decreased at this time point. However, the IL8 protein secretion was increased at 72 hrs. In parallel, IL6 protein secretion was enhanced for both 24 and 72 hrs post UCB treatment, while mRNA expression showed 1.5-fold increase only at 72 hrs. *TNF-α* levels did not show any elevation upon UCB exposure for the UM51 line (0.7 and 0.8-fold at 24 and 72 hrs), but slightly increased in both secretome and mRNA expression with 1.5-fold change in the i705-C2 line. Fernandes et al., previously reported that UCB enhances TNFR1 protein level in neural cells (such as astrocytes) along with a time-dependent release of multiple cytokines such as TNF-α, IL-1ß and IL-6. But neurons secrete merely mild levels of IL-6, even to a less extent TNF-α, and almost undetectable amounts of IL-1ß upon UCB exposure [28]. The correlation of mRNA and secretome expression observed in our results suggest that mRNA and protein expression might have a slight time shift in their expression as a response to stress to induce inflammation in CNS. For example, both of cell lines showed increased IL1-ß secretion only at 24 hrs, while IL1-ra secretion was increased merely at 72 hrs. These observations point to a variation in the levels and patterns of initiation of inflammation post UCB treatment. Secretome analysis also revealed enhanced vascular endothelial growth factor (VEGF) secretion and reduced sex hormone-binding globulin (SHBG) secretion upon UCB treatment for all the conditions, except UM51 24hrs. Previous studies described low VEGF expression in adult human brain and upregulated VEGF levels has been observed in brain injury or in chronic neuro-inflammation [40,41]. Increased *VEGF* expression might be a cause or response to bilirubin treatment, which remains a question because of the multiple roles that VEGF plays. On the other hand, SHBG exhibits anti-inflammatory effects in macrophages and adipocytes, however, SHBG expression in brain is not yet known [42]. A slight reduction of *SHBG* expression with 0.6-fold (UM51) and 0.8-fold (i705-C2) at 72hrs post UCB treatment might indicate a tendency towards initiation of anti-inflammatory response, while VEGF expression was increased at this time point. Additionally, enhanced secretion of other pro-inflammatory cytokines such as IL1-ß, IL16, IL1-ra, IL-11, INF-G, were observed along with anti-inflammatory cytokine IL-13 secretion. The secretome profile unveiled a pro-inflammatory CNS environment with variable cytokine secretion for both of the healthy and disease cell lines at different time points, thus indicating pronounced neuro-inflammation upon UCB treatment. Overall, in accord with Brites et al., the increase in mRNA expression and secreted levels of *IL6, IL8 and TNF-α* along with other inflammatory cytokines points to the onset or initiation of neuro-inflammation in the cortical organoids upon UCB treatment [3,28,43].

Next, we performed RNA microarray-based global gene expression analysis to have an overview of the molecular effects of UCB on cortical organoids such as differential gene expression and associated biological processes (GO-BP) and KEGG pathways for both treatment time points for the cell lines. Both cell lines showed activation of cytokine-cytokine receptor interaction, calcium-signaling pathway, MAPK signaling pathway, and neuroactive ligand receptor interaction among their uniquely expressed gene sets. In the patient line, cytokine-cytokine receptor activation was activated for both 24 and 72 hrs in the UCB treated condition, whereas in the healthy line this pathway seemed to be activated at 72 hrs. The observed GO terms and KEGG associated pathways did not show activation of any of these mentioned pathways at th 24 hrs treated condition in healthy UM51 line. Activation of the GO-BP-inflammatory bowel disease was observed in the i705-C2-24 hrs and UM51-72 hrs post treated conditions. These findings point towards a possibility that the patient line shows an accelerated response to initiate inflammatory responses (24 hrs in this case) than the healthy line. Slightly enhanced mRNA expression of *TNSF12* (1.12-fold in UM51, 1.6-fold in i705-C2), *AZI2* (1.2-fold in patient line), *MyD88* (1.2-fold in patient line) were observed upon UCB treatment. TNFSF12 can induce pro-inflammatory cytokines such as IL-6, MCP-1, IL-8, and MMP-9 [44]. Myeloid differentiation primary response factor 88 (MyD88) is an intracellular adapter protein. Many of the Toll like receptors (TLRs) and cytokine receptors signal through MyD88 and co-ordinate the mediation of pro-inflammatory signaling cascades [45-47]. 5-azacytidine-induced protein 2 (AZI2) is another adaptor protein and contributes to the activation of NFκB dependent gene expression. AZI2 belongs to TNFR family and associated with NFκB activator (TANK)–binding kinase 1 (TBK1) binding protein family [47,48]. Increased expression of these genes’ points to the initiation of inflammation upon UCB treatment.

GO-BP terms revealed at 72 hrs the activation of positive regulation of NFκB transcriptor factor activity (*AR, TRADD, RPS6KA4*) in i705-C2 line, while positive regulation of CREB transcription factor activity (*RELN, RPS6KA4*) and NLRP3 inflammasome complex assembly (*GBP5, NLRP3*) in the UM51 line. *NLRP3* is an upstream activator of NFkß signaling. The activation and increased NLRP3 mRNA expression might indicate NFκB mediated inflammatory responses upon UCB exposure. Similar to astrocytes and microglia, UCB induced NFκB activation was observed in neurons as well, but at lower levels [11,28]. Based on the cellular context, NFκB plays diverse roles in the CNS. Axonal growth of neurons can be implicated with NFκB activation [49,50]. NFκB may also regulate neural development, plasticity, and neurogenesis, when it is activated in the synapse and transported to the cell nucleus [28,51,52]. From our observed results, GO-BP terms of the common set of genes revealed neurodevelopmental pathways to be upregulated for both cell lines at 72 hrs post UCB treatment with the activation of cellular developmental process, nervous system development, axon development, axon guidance, tight junction, positive regulation of axogenesis, positive regulation of synapse assembly.

On the other hand, regulation of MAPK cascade was also observed in both UM51 and i705-C2 lines upon UCB treatment. MAPK signaling pathways were observed to be activated in UCB-treated astrocytes as well [28]. A slight enhancement of phospho-P38 protein expression with 1.25-fold increase was observed for the patient line post UCB exposure by Western blot analysis.

KEGG pathways associated with the common set of genes revealed DNA damage and repair -related pathways such as P53 signaling pathway, homologous recombination, Fanconi anemia pathway as upregulated in the UM51 line at 24 hrs. These pathways were downregulated at 24 hrs in the i705-C2 line. With respect to DNA damage, P53 induces multiple classes of its target genes, such as metabolic genes, DNA repair genes, cell-cycle arrest, and cell death effectors [53]. The UM51 line showed enhanced mRNA expression of DNA damage-repair and apoptosis related genes such as *P53, BCl2, ATM, ATR, CHEK1*, and *CHEK2* compared to the i705-C2 line. P53 protein expression was 1.64-fold upregulated at 24 hrs in the healthy line and the level decreased at 72 hrs. Conversely, there was no increase in P53 protein expression (1.1-fold at 72 hrs) in the patient line, but an increase of *P53* mRNA expression (1.6-fold) was observed at 72 hrs. P53 signaling and DNA damage repair-related pathways appeared to be activated at 24 hrs in the healthy line. Interestingly, a number of inflammatory and development-related pathways (neuroactive-ligand receptor interaction, cAMP signaling pathway, cytokine-cytokine receptor interaction) were downregulated at 24 hrs in the UM51 line. Indications of inflammatory responses were observed at 72 hrs, implying a delayed initiation of inflammation in the healthy line. In parallel, at 24 hrs, the i705-C2 line showed activation of NFkß, PI3K, and chemokine signaling pathways thus implying the initiation of inflammation, whilst cellular developmental processes such as Notch and TGFß signaling pathways were upregulated. Expression of *NLRP3* which is known to play a role in inflammation, immune responses, and apoptosis was 1.9-fold enhanced in the i705-C2 line only [54].

On the other hand, positive regulation of CREB transcription factor activity could be a response to stress or induction of inflammatory cascades [55]. Zhang et al., described that UCB can regulate Ca2+ channel opening [56]. This might have a correlation with the observed activation of calcium signaling pathway upon UCB exposure in the KEGG pathway analysis. Ca 2+ influx into cortical neurons regulate the expression of nNOS mediated by a CREB family transcription factor [56]. The positive regulation of CREB transcription factor expression levels was observed in the UM51 cell line. Of note, CREB transcription factors are associated with inflammation and apoptotic cell death, while both of these could be possible effects of UCB on the neural organoids [57]. The control line-UM51 showed 1.2-fold increase in both mRNA and protein expression of CREB and 1.2-fold increase of phospho-CREB protein expression. Apoptotic cell death was observed in both cell lines at 24 hrs post UCB exposure. However, the UM51 line showed more than 30% increase in apoptotic cell death, while the patient line showed around 20% increase. γH2AX, which can be a marker for both DNA damage and apoptotic cell death, also underwent a 1.4-fold increase in protein expression in the UM51 line [58].

Through all the observed analyses, the UM51 line showed increased apoptotic cell death and DNA damage and repair-related gene expression at 24 hrs post UCB treatment, and then the activation of inflammatory related pathways at 72 hrs. On the other hand, the i705-C2 line did not show increased DNA damage repair-related gene expression at 24 hrs (*ATM, ATR, CHEK1, CHEK2*). However, the i705-C2 line leaned towards initiation of inflammation at 24 hrs, which then decreased at 72 hrs. It can be assumed from these observations-distinct activated pathways and the associated gene expressions, are indicative of a switch or shift in initiation of inflammation and cell death related pathways in these cell lines after treating with UCB. Although both cell lines showed inflammatory responses after UCB treatment, they seemed to be adopting distinct pathways and time point to respond to the UCB-induced stress which results in apoptotic cell death, DNA damage repair, or inflammation.

## 5. Conclusion

In summary, this study provides valuable molecular insights into UCB-induced neuro-inflammation in iPSC-derived cortical organoids. The global gene expression analyses provided an overview of distinct pathways and genes, which might be associated with the neuro-inflammatory effects caused by UCB. The iPSCs-derived cortical organoids employed in this study represent a Crigler-Najjar syndrome model to study *UGT1A1* mutation and its subsequent phenotypic manifestation and potential application for future BIND associated toxicological studies and drug screening.

## Supplementary Materials

The supplementary materials include: Table S1: Composition of Cortical organoid differentiation medium, Table S2: List of utilized antibodies, Table S3: List of used qRT-PCR primers, Figure S1: Generation of UM51 and i705-C2 iPSC-derived HLCs to check the presence or expression of UGT1A1, Figure S2: The secretome analysis of each individual sample using Human XL Cytokine Array, Figure S3: Selected GO terms with corresponding genes and KEGG pathways from the uniquely expressed genes post UCB exposure, Figure S4: WB analysis and Dot plots of upregulated GO.BP terms from the common set of genes at 72hr post UCB treatment, Figure S5: Immunofluorescene staining showing the expression of γH2AX at 24hrs post UCB exposure. Supplementary File S1: Complete lists of KEGG pathways with the associated unique gene sets of bilirubin treated i705-C2 and UM51 organoids at 24 and 72 hrs. Supplementary File S2: Complete lists of up- and down-regulated KEGG pathways in the common gene-set of i705-C2 and UM51 control and bilirubin treated condition at 24 hrs. Supplementary File S3: Complete lists of GO terms based on uniquely expressed gene-sets of bilirubin treated i705-C2 and UM51 cortical organoids at 24 and 72 hrs. Supplementary File S4: Complete list of GO terms based on the upregulated common gene-set between i705-C2 and UM51 control and bilirubin treated condition at 72 hrs. Supplementary File S5: All the whole blot membranes that have been cropped and depicted in the main and supplementary figures.

## Author Contribution

A.I.P. designed and performed experiments, processed, and analysed the data, wrote and edited the manuscript. W.W. performed the bioinformatic analysis, data curation, helped with the figures, wrote the bioinformatic section in the methods and materials, and edited the manuscript. J.A. conceptualized and designed the work, edited the manuscript, acquired funding, and supervised the study. All authors have read and agree to the submitted version of the manuscript.

## Acknowledgement

J.A. acknowledges the medical faculty of Heinrich Heine University for financial support.

## Institutional Review Board Statement

The study was conducted according to the guidelines by the Ethics Committee of the medical faculty of Heinrich-Heine University, Germany (protocol code: 5704).

## Data availability statement

All microarray data will be available at NCBI GEO server when the manuscript is accepted.

