## Supplementary Figure Legend for "Unconjugated bilirubin induces neuro-inflammation in an induced pluripotent stem cell-derived cortical organoid model of Crigler Najjar Syndrome"

**Supplementary Figure 1: Generation of UM51 and i705-C2 iPSC-derived HLCs to check the presence or expression of UGT1A1.**

**(a)** (i.ii), **(b).** (i,ii) IF showed characterization of the iPSC-derived HLCs by using typical HLC markers. Both cell lines showed HNF4a, E-cadherin, CYP2D6, AFP and albumin protein expression. The UGT1A1 was only expressed on the healthy line. Scale bars depict 50 µm. **(c)** CYP3A4 and CYP2D6 assays were performed to check the functionality of the generated HLCs. **(d)** qPCR analysis showed reduced but presence of *UGT1A1* expression in mRNA level along with expression of other HLC markers, such as *AFP, albumin* and *CYP2D6*. Depicted values are mean of three different independent (n=3) experiments. Error bar depicts ± 95% confidence interval. Values were normalized to RPL0 (housekeeping gene) and subsequently to DMSO treated control organoids. **(e)** (i,ii) A very fade expression of UGT1A1 was observed in WB analysis. HEPG2 was used as a positive control. The bar graph represents the UGT1A1 expression in fold change. Values were normalized to GAPDH (housekeeping gene).

**Supplementary Figure 2: The secretome analysis of each individual sample using Human XL Cytokine Array.**

**(a)** Individual image of UCB treated and non-treated Human XL cytokine array membranes for both cell lines and time points. **(b)** The complete heatmap of each condition, representing the secretome profile, which was obtained after analysis the membrane dots. **(c)** mRNA expression of *VEGFA* (i) and *SHBG* (ii) after UCB treatment. Depicted values are mean of three different independent (n=3) experiments. Error bar depicts ± 95% confidence interval. Values were normalized to RPL0 (housekeeping gene) and subsequently to DMSO treated control organoids.

**Supplementary Figure 3: Selected GO terms with corresponding genes and KEGG pathways from the uniquely expressed genes post UCB exposure.**

**(a)** Selected GO terms showing the corresponding genes from the uniquely expressed genes after UCB treatment on both cell lines at 72 hrs time point. **(b)** qPCR analysis showing a slight increase in mRNA expression of *NLRP3* (1.9-fold at UM51 24hrs),*CREB1* (1.15-fold at UM51 72hrs, i705-C2 24hrs) , *TNFSF12* (1.5-fold at i705-C2 72hrs), *AZI2* (1.2-fold at i705-C2 24hrs), and *MyD88* (1.2-fold at i705-C2 72hrs) for certain time points. These genes were selected from the mentioned GO terms. Depicted values are mean of three different independent (n=3) experiments. Error bar depicts ± 95% confidence interval. Values were normalized to RPL0 (housekeeping gene) and subsequently to DMSO treated control organoids. **(c)** Dot plot from KEGG associated pathways for healthy cell line, did not show an initiation of inflammatory related pathways at 24hrs.

**Supplementary Figure 4: WB analysis and Dotplots of upregulated GO.BP terms from the common set of genes at 72hr post UCB treatment.**

**(a). (b). (c)**: WB analysis showing CREB, Phospho-CREB and phospho-P38MAPK protein expression. **(a)** (i, ii) **(b)** (i, ii) The UM51 showed slight increase in protein expression for both CREB and phosphor-CREB at 24 hrs, while the i705-C2 line did not show an increase. **(c)** (i) Slightly increased phospho-P38MAPK expression was observed in the patient line for both 24 and 72 hrs time point. **(a)** (iii), **(b)** (iii), **(c)** (ii), Bar graphs showing the protein expression in fold change. Values were normalized to ß-Actin **(a), (b)** and GAPDH **(c)** (housekeeping gene) and subsequently to DMSO treated control organoids. **(d)** (i, ii) GO terms of the common set of genes showing the neurodevelopmental pathways to be upregulated for both cell lines at 72 hrs post UCB treatment.

**Supplementary Figure 5: Immunofluorescene staining showing the expression of γH2AX at 24hrs post UCB exposure.**

**(a), (b)** 24 hours post treatment, in both cell lines IF showed a slight increase in the presence of DNA damage and repair related and apoptotic marker y-H2AX compared to the control, while proliferation marker Ki-67 did not show a difference comparing to the control (i,ii). Scale bars depict 100 µm. Confocal imaging of the Y-H2AX (green), Ki67 (red) positive cells showing a higher magnification. Scale bars depict 20µm.
