## Supplementary Table S1 for "Unconjugated bilirubin induces neuro-inflammation in an induced pluripotent stem cell-derived cortical organoid model of Crigler Najjar Syndrome"

*Table S 1: Composition of Cortical organoid differentiation medium.*

| Component | % |
|---|---|
| Neurobasal medium | 50 |
| DMEM-F-12 | 50 |
| Glutamax | 1 |
| MEM NEAA | 1 |
| Pen/strep | 1 |
| ß-Mercaptoethanol | 0.1 |
| N2 | 0.5 |
| B27 (without Vit A) | 1 |

| | |
|---|---|
| Insulin | 23µM |
| SB431542 | 5µM |
| Dorsomorphin | 0.5µM |
