## Supplementary Table S2 for "Unconjugated bilirubin induces neuro-inflammation in an induced pluripotent stem cell-derived cortical organoid model of Crigler Najjar Syndrome"

*Table S2: List of utilized antibodies.*

| Primary antibody | Application | Dilution |
|---|---|---|
| Anti-rabbit SOX-2 | IF | 1:100 |
| Anti-mouse ß3 tubulin (TUJ1) | IF | 1:250 |
| Anti-rabbit MAP2 | IF | 1:250 |
| Anti-guinea pig DCX | IF | 1:200 |
| Anti-rabbit cleaved caspase 3 | IF | 1:800 |
| Anti-mouse P53 | WB | 1:1000 |
| Anti-mouse γH2A.X | IF, WB | 1:200, 1:50 |
| Anti-rabbit UGT1A1 | IF, WB | 1:200, 1:1000 |
| Anti-rabbit HNF4α | IF | 1:250 |
| Anti-rabbit CYP2D6 | IF | 1:200 |
| Anti-rabbit AFP | IF | 1:200 |
| Anti-mouse Albumin | IF | 1:500 |
| Anti-rabbit ECAD | IF | 1:100 |
| Anti-mouse KI67 | IF | 1:200 |
| Anti-rabbit CREB | WB | 1:1000 |
| Anti-rabbit Phospho CREB | WB | 1:1000 |
| Anti-rabbit Phospho-P38MAPK | WB | 1:1000 |
| Anti-mouse ß-Actin | WB | 1:5000 |
| Anti-mouse GAPDH | WB | 1:1000 |

| Secondary antibody | Application | Dilution |
|---|---|---|
| Alexa 488 goat anti rabbit IgG (H+L) | IF | 1:600 |
| Alexa 555 goat anti mouse IgG (H+L) | IF | 1:600 |
| Alexa 647 goat anti guinea pig | IF | 1:600 |
| Anti-rabbit IgG, HRP-linked Antibody | WB | 1:1000 |
| ECL peroxidase labelled anti mouse Antibody | WB | 1:5000 |

| Company | Order no |
|---|---|
| Cell Signaling | 3579S |
| Cell Signaling | 4466S |
| SySy | 188002 |
| Millipore | AB2253 |
| CST | 9664S |
| CST | 9283S |
| CST | 9718S |
| Thermo Fisher | PA5-98229 |
| Abcam | 92378 |
| Abcam | ab185625 |
| Sigma | HPA023600 |
| Sigma | A6684-.2ml |
| CST | 3195 |
| CST | 9449S |
| CST | 9197S |
| CST | 9198S |
| CST | 9212S |
| CST | 3700S |
| Ambion/Invitrogen | AM4300 |

| Company | Order no |
|---|---|
| ThermoFisher | A11008 |
| ThermoFisher | A21424 |
| ThermoFisher | A21450 |
| CST | 7074S |
| GE | NA931V |
