## Supplementary Table S3 for "Unconjugated bilirubin induces neuro-inflammation in an induced pluripotent stem cell-derived cortical organoid model of Crigler Najjar Syndrome"

*Table S3: List of used qRT-PCR primers.*

| Primer | Forward 5'→3' | Reverse 5'→3' |
|---|---|---|
| IL-6 | GGTACATCCTCGACGGCATCT | GTGCCTCTT TGCTGCTTTCAC |
| IL-8 | GTGCAGTTTTGCCAAGGAGT | ACTTCTCCACAACCCTCTGC |
| TNF- α | AGAACTCACTGGGGCCTACA | AGGAAGGCCTAAGGTCCACT |
| P53 | CAGGGCAGCTACGGTTTCC | CAGTTGGCAAAACATCTTGTTGAG |
| BCL2 | CTTCAGGGATGGGGTGAACT | CAGCCTCCGTTATCCTGGAT |
| MDM2 | AAACTGGGGAGTCTTGAGGG | TGCACATTTGCCTGCTCCTC |
| ATM | ACACTGCGCGTATAAGCCAATC | TTTTCAACCAGTTTTCCGTTACTTC |
| ATR | TGTTGGGCCCACTTTATGCAGC | TAGAGACGACCTGAGACGACGC |
| CHEK1 | GGATCAGCTTTTCCCAGCCCAC | TTCTGTGAGGATCCTGGGGTGC |
| CHEK2 | ACGGAGTTCACAACACAGCAGC | TCTACTAGTCGAAAGCGGCCCC |
| UGT1A1 | AACTGCCTTCACCAAAATCCACT | TTGCCATAGCTTTCTTCTCTGG |
| VEGF | AAGGAGGAGGGCAGAATCATC | TGATGTTGGACTCCTCAGTG |
| SHBG | GGCTGGATGATGGGAGATGG | ATCTCGGCCTGTTTGTCCAG |
| NLRP3 | TGTGAAACGCTCCAGCATCC | CAACATGCTGATGTGAGGCA |
| CREB1 | GGAATCTGGAGCCGAGAACC | TGGACTTGAACTGTCTGCCC |
| TNFSF12 | TAAAGGCCGGAAAACACGGG | GACAGCCTTCCCCTCATCAAA |
| AZI2 | CTGGTGACTCAAGTACAAGCTG | CTTCACTGCATCCTACAGGGG |
| MyD88 | CCACACTTGATGACCCCCTG | ATCAGAGACAACCACCACCA |
| RPL0 | TCGACAATGGCAGCATCTAC | ATCCGTCTCCACAGACAAGG |
