## Supplementary File 5 for "Unconjugated bilirubin induces neuro-inflammation in an induced pluripotent stem cell-derived cortical organoid model of Crigler Najjar Syndrome"

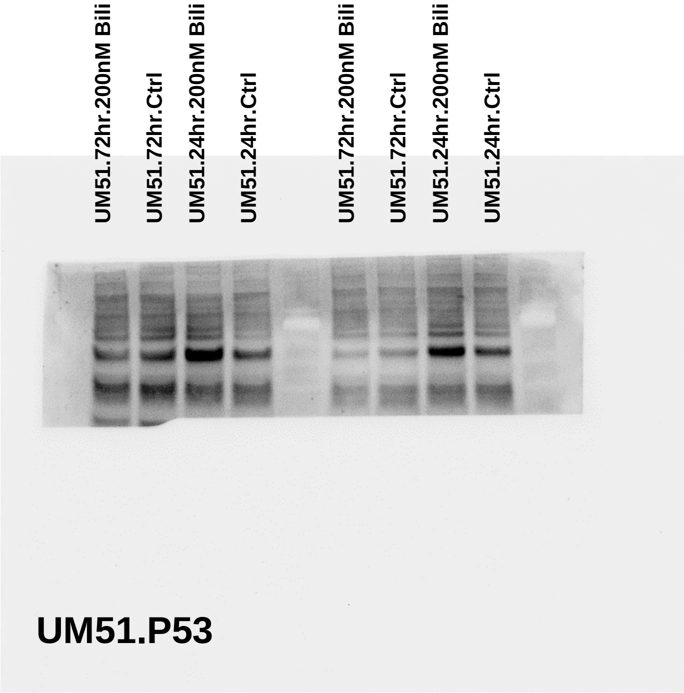


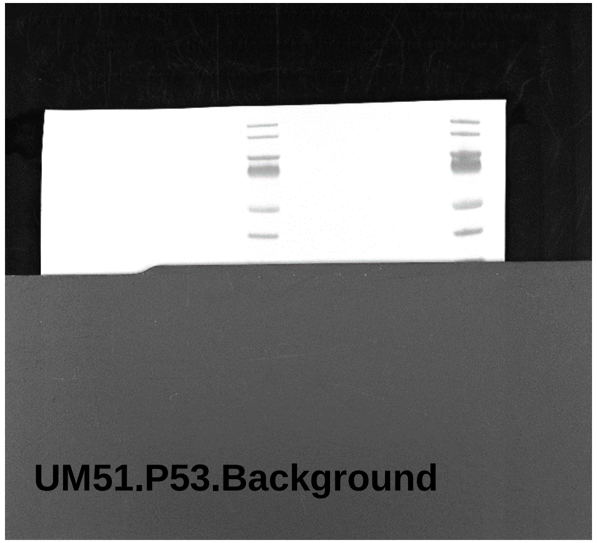


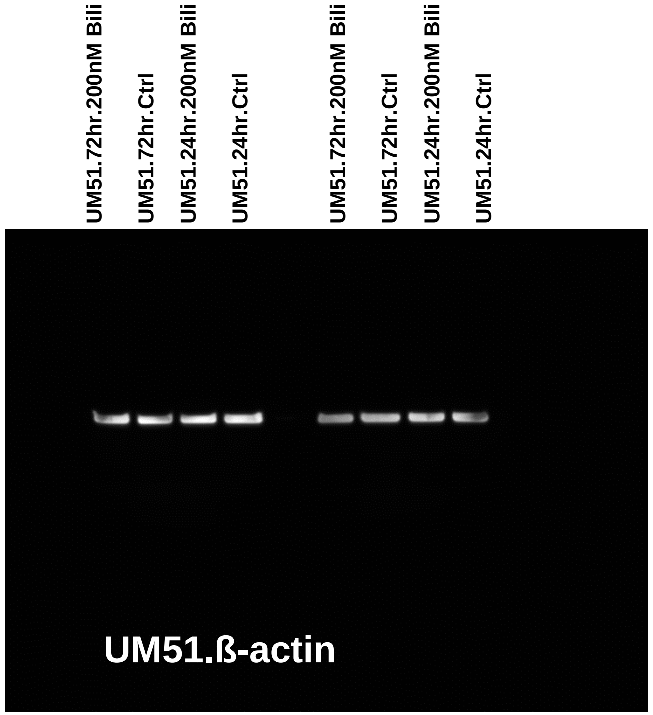


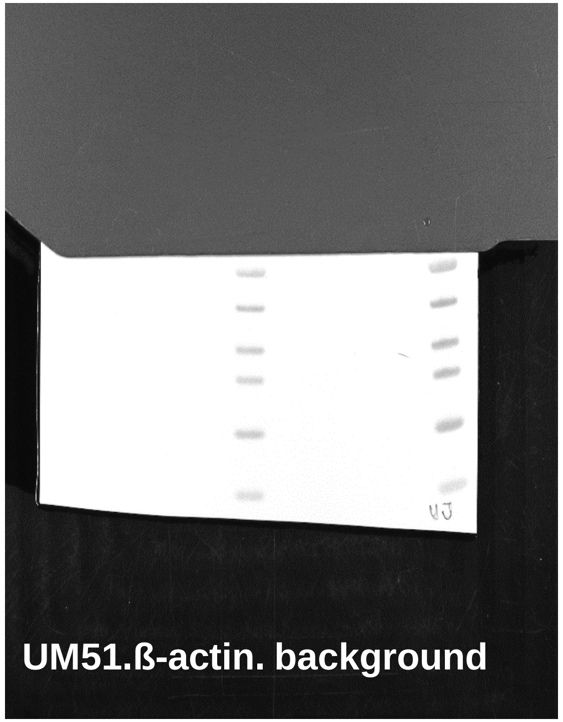


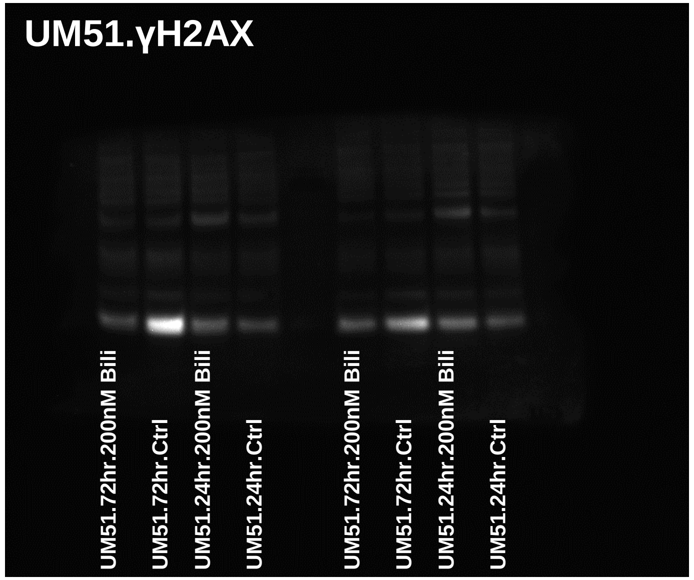


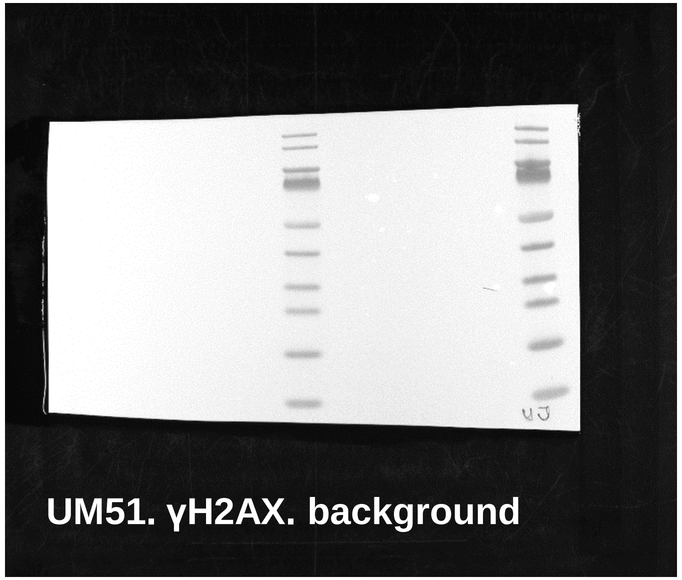


The whole blot membranes for main figure 4, 5. The stainings were done on the same membrane for P53 and γH2AX.

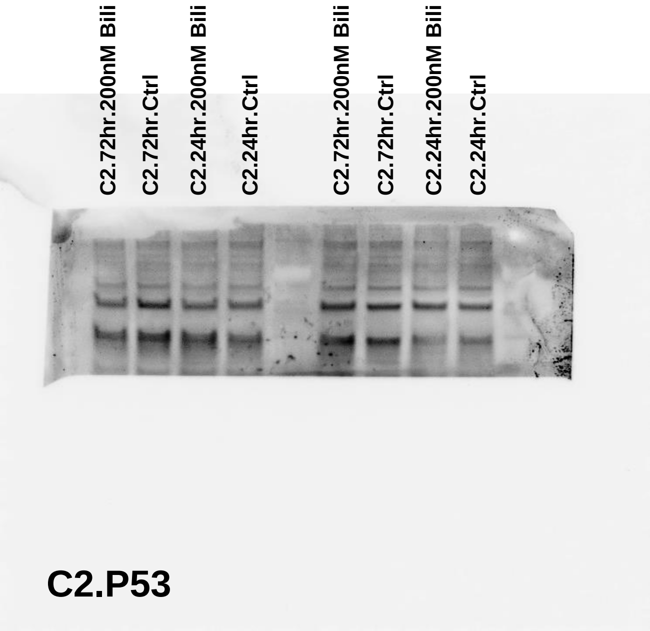


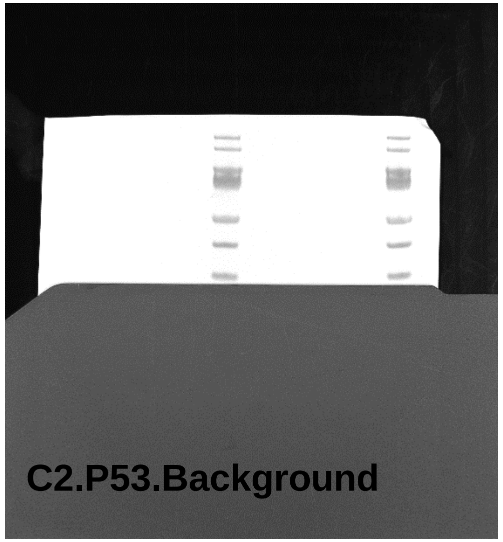


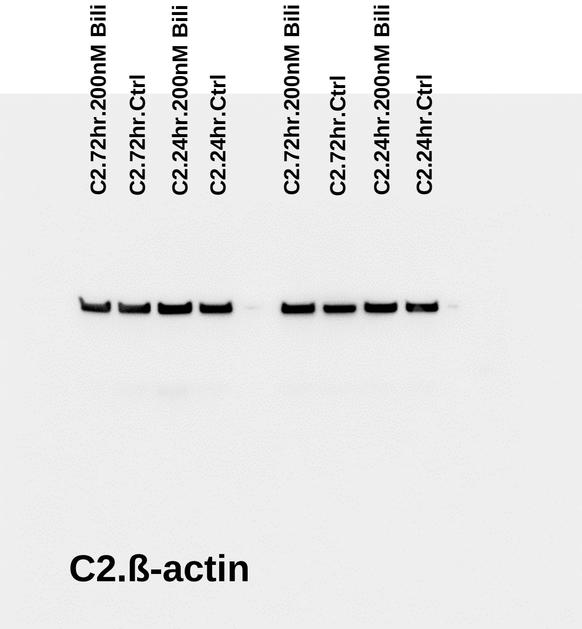


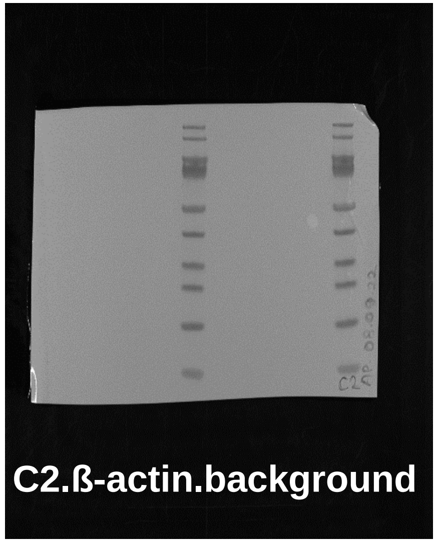


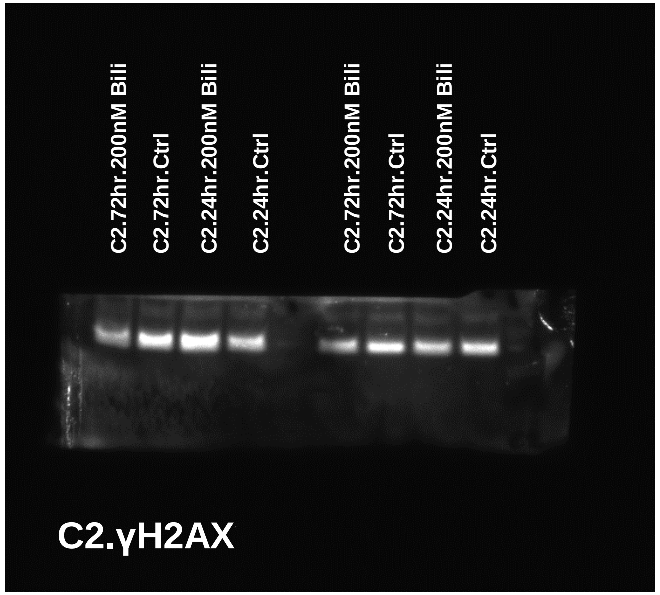


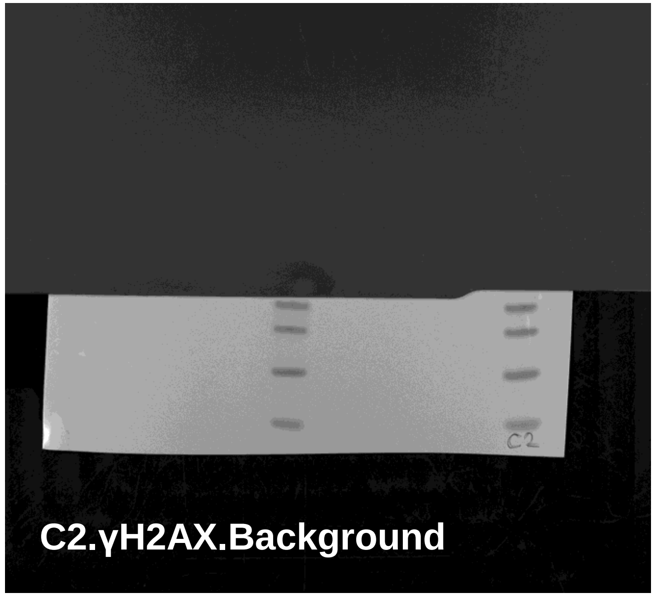


The whole blot membranes for main figure 4, 5. The stainings were done on the same membrane for P53 and γH2AX.

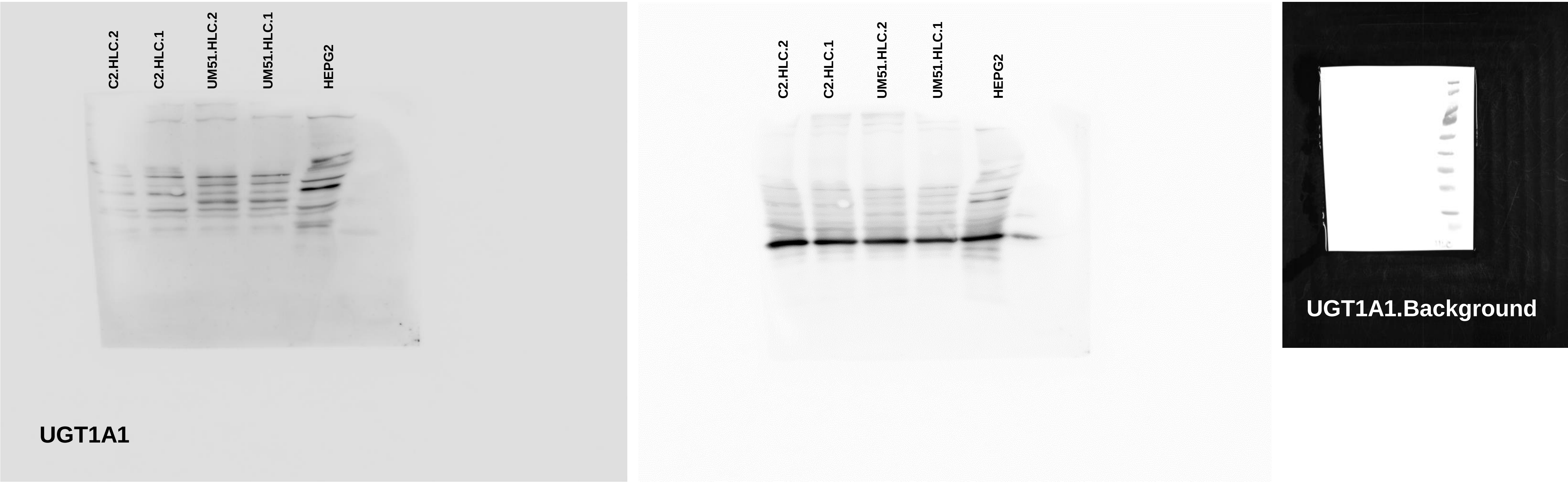


The whole blot membranes for supplementary figure 1.

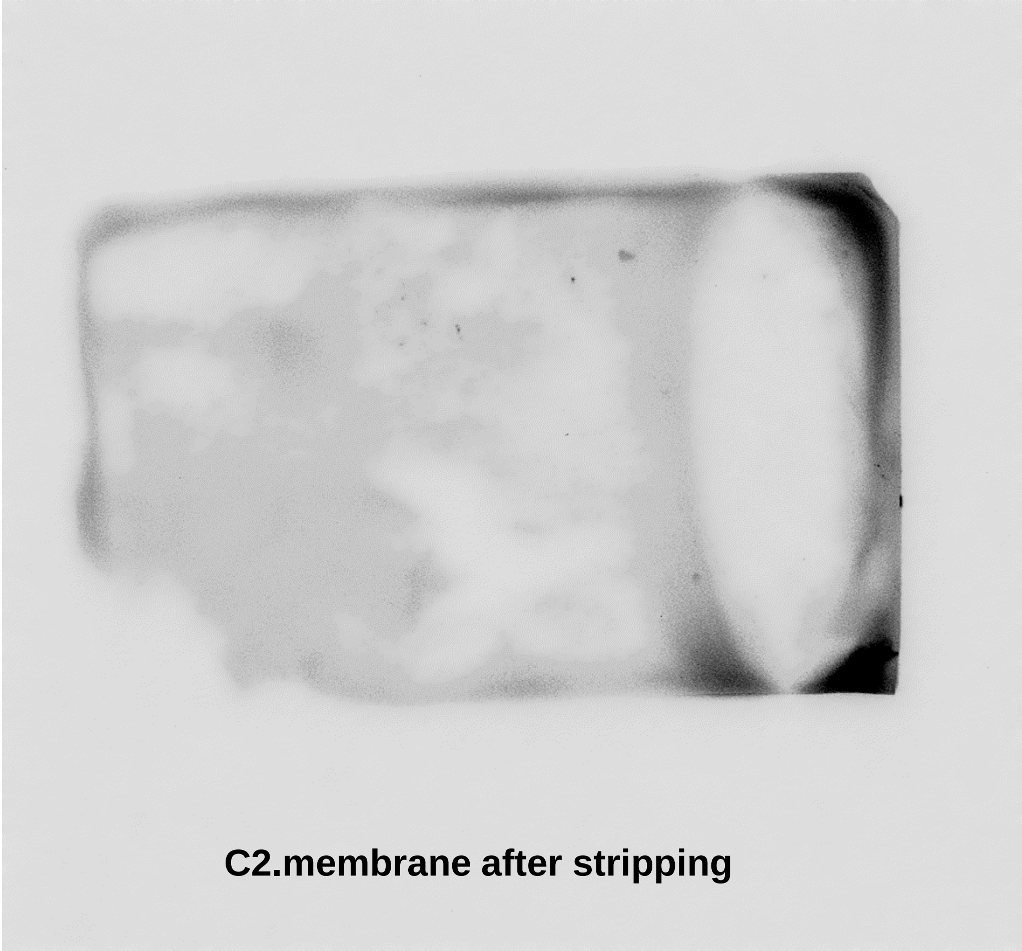


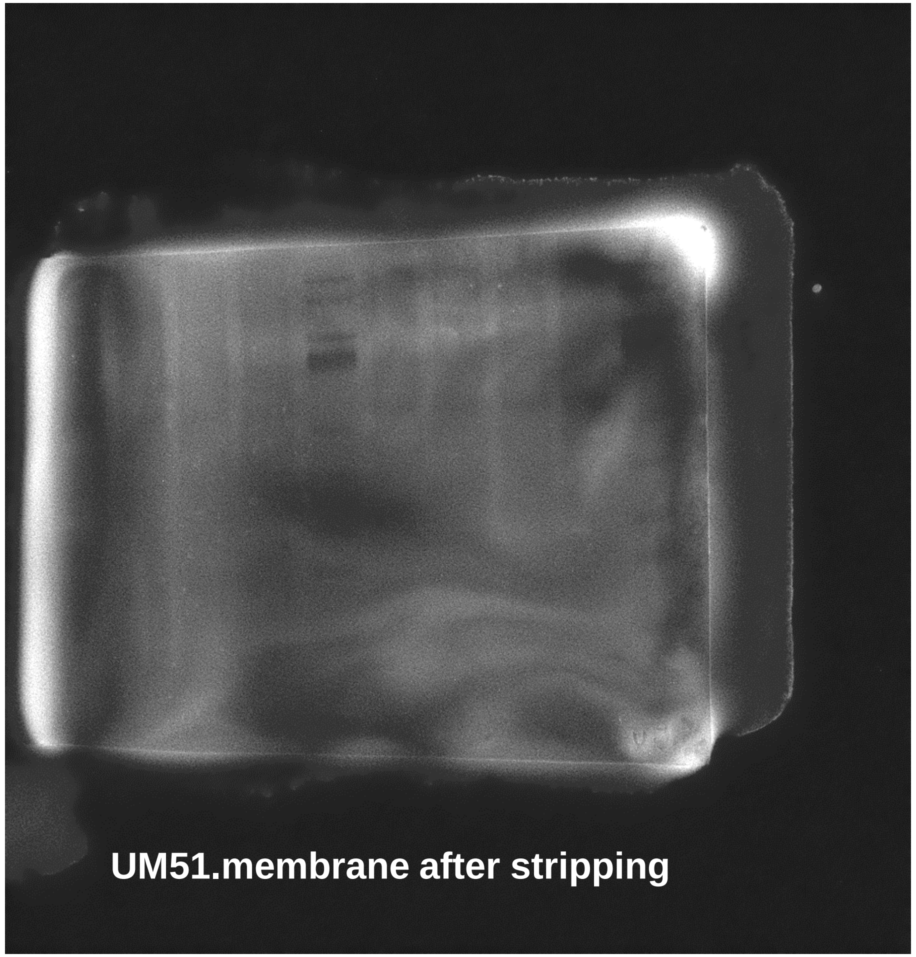


The blot membranes which were used on main figure 4,5 were stripped and used for the staining of supplementary figure 4a, 4b.

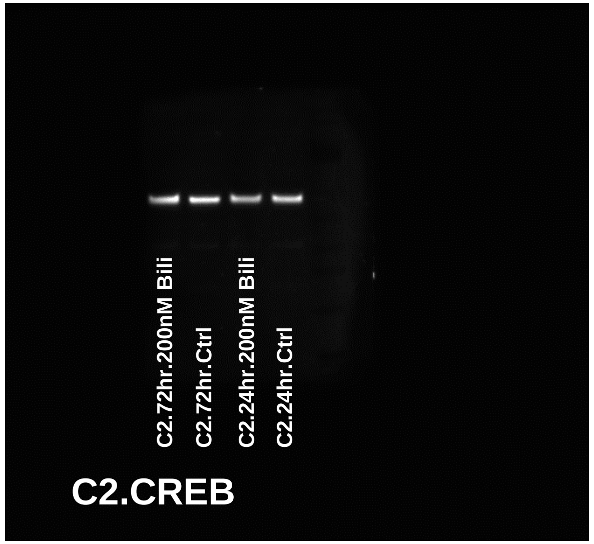


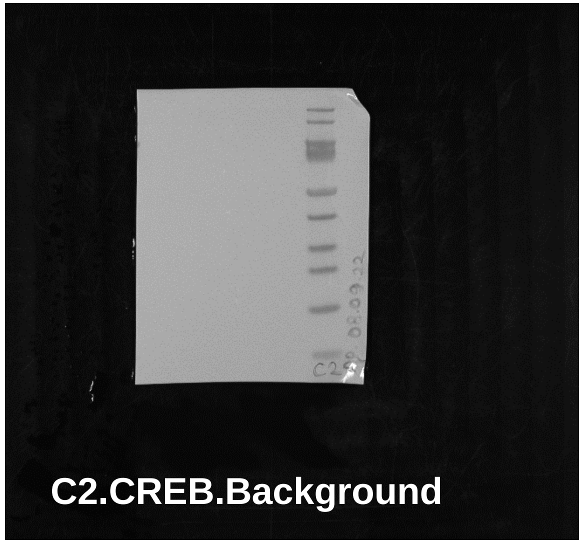


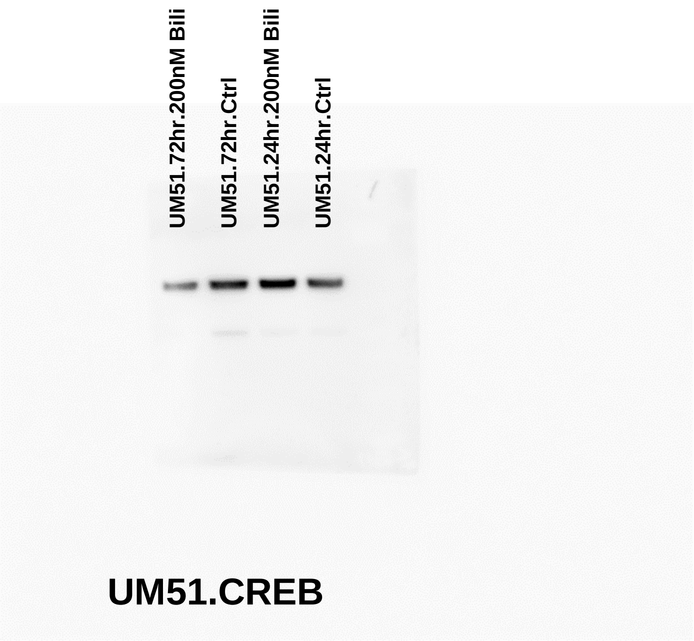


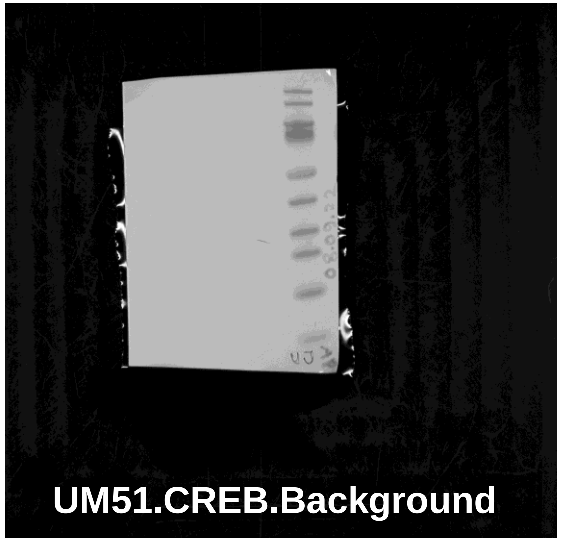


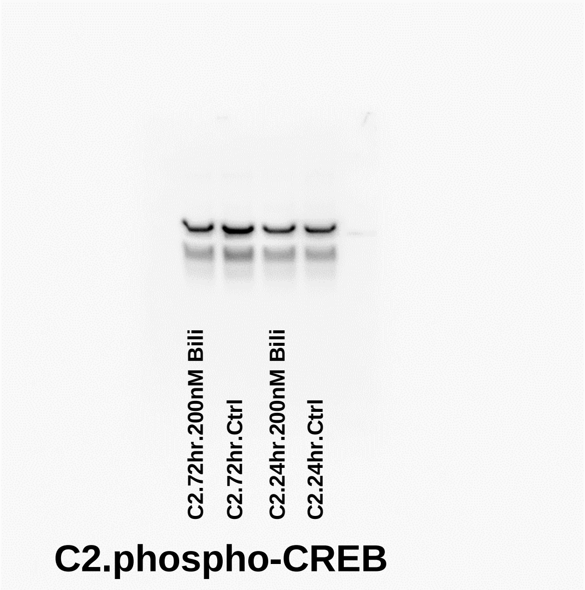


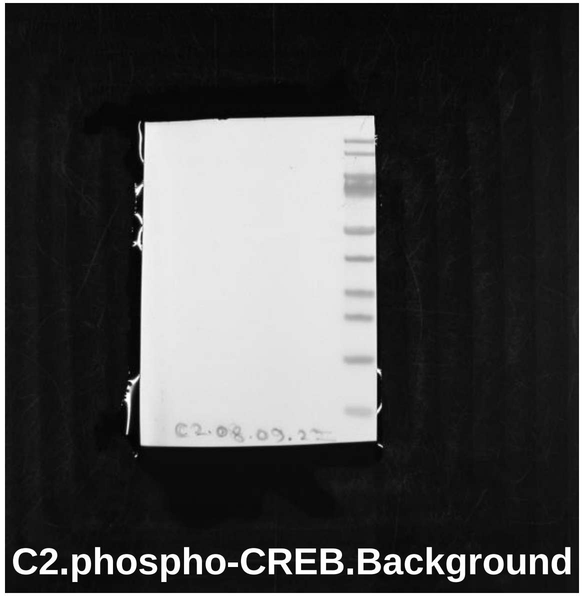


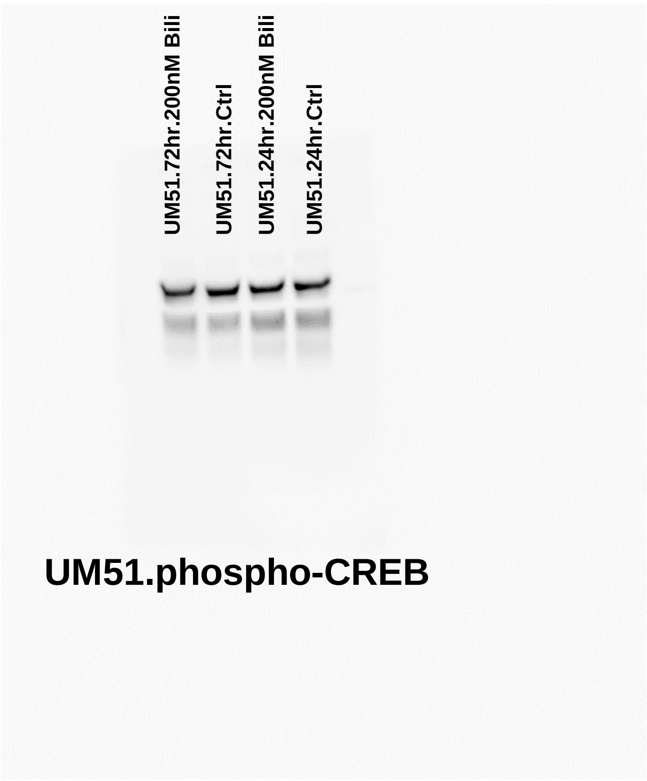


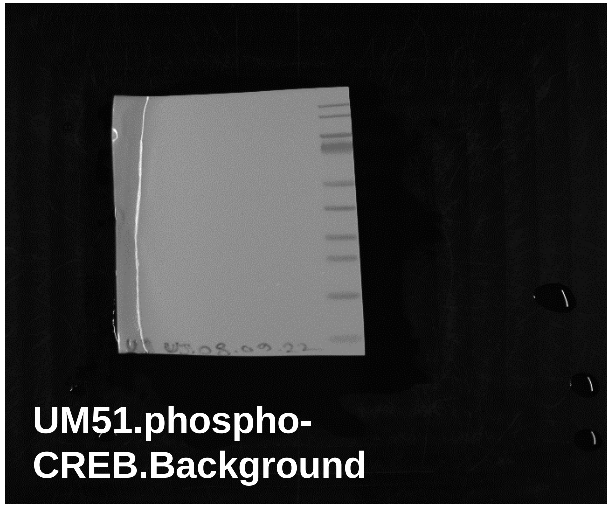


The whole blot membranes for supplementary figure 4a, 4b.

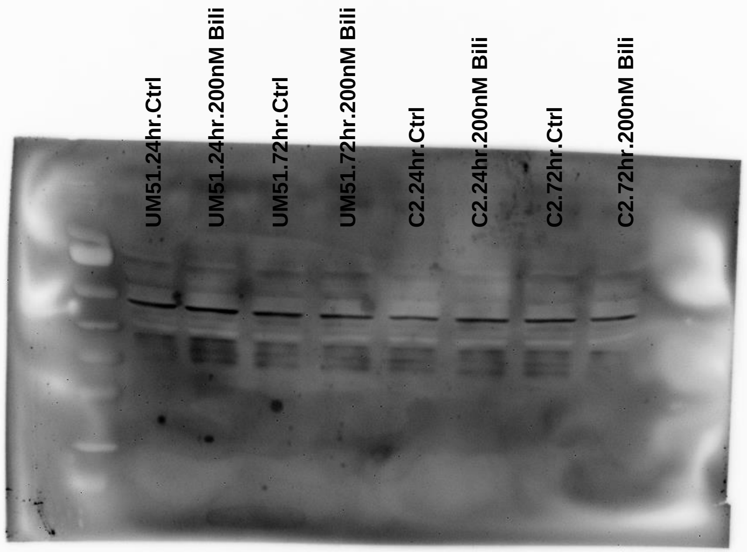


**Phospho-p38MAPK**

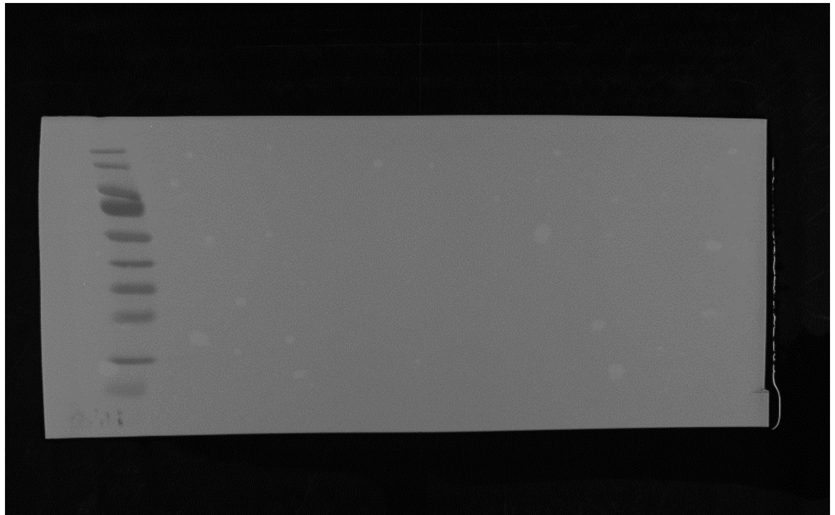

**Whole membrane for phospho-p38MAPK**

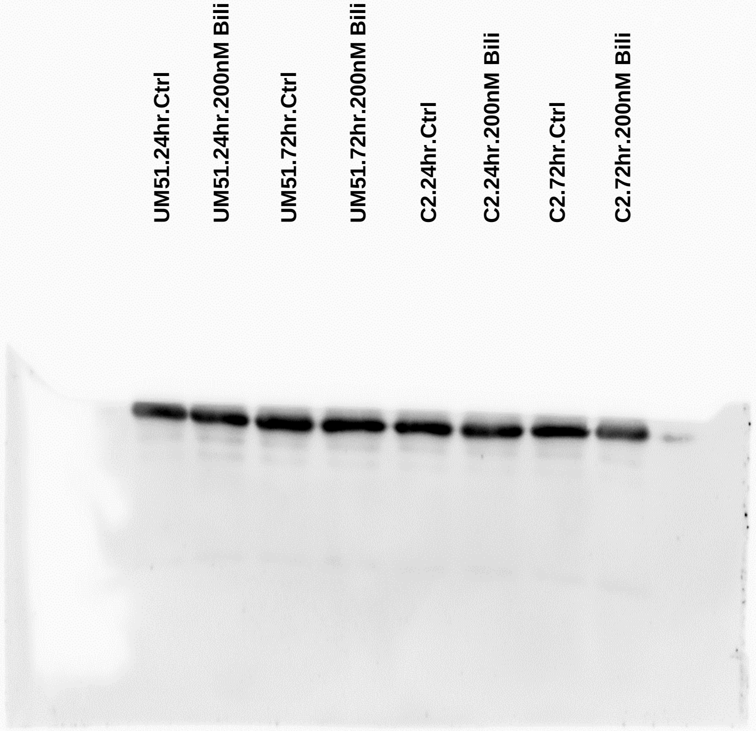


**GAPDH**

The whole blot membranes for supplementary figure 4c.
